# Breaking the link between demyelination and axon loss: SARM1 inhibition as a neuroprotective strategy for multiple sclerosis

**DOI:** 10.64898/2026.09.25.754536

**Authors:** Micah Feri, Stephanie R. Peterson, Melika Rezanejad, Flavio Denzel Cardenas, Alyssa M. Anderson, Brandon T. Poole, Moyinoluwa T. Ajayi, Sung Hoon Kim, John K. Katzenellenbogen, Seema K. Tiwari-Woodruff

## Abstract

Axonal degeneration is a principal driver of irreversible neurological disability in multiple sclerosis (MS), yet current treatments fail to target this neurodegenerative phase directly. Sterile alpha and TIR domain-containing protein 1 (SARM1) has emerged as an executioner of programmed axon destruction and is overexpressed in grey and white matter MS tissue, making its inhibition a promising therapeutic strategy. However, the effects of SARM1 inhibition on localized neurodegeneration versus systemic and central inflammation remain poorly understood in complex autoimmune environments like MS. Here, we investigated SARM1 pathology using global SARM1-/- knockout (SARM1-/-) mice, AAV-mediated CRISPR knockdown and overexpression of SARM1 in retinal ganglion cells (RGCs), and the small-molecule SARM1 inhibitor 5-iodoisoquinoline (5IIQ), across optic nerve crush (ONC) and experimental autoimmune encephalomyelitis (EAE) models. While global SARM1-/- did not alter the overall clinical course of EAE, it revealed a complex phenotype characterized by an altered peripheral inflammatory cytokine profile and persistent CNS immune infiltration, alongside preserved axonal and myelin integrity. RGC-restricted SARM1 knockdown partially preserved RGCs and axons during EAE, whereas SARM1 overexpression worsened both retinal function and axonal injury. Pharmacological inhibition with 5IIQ preserved axonal integrity and restored visual function in both the ONC and EAE models, as confirmed by electrophysiology, and reduced serum neurofilament light chain (NfL) in EAE, without affecting demyelination. Together, these findings demonstrate that SARM1 inhibition uncouples axonal self-destruction from demyelination and gross neuroinflammation while providing robust structural and functional neuroprotection. This study establishes SARM1 as a viable target for neuroprotective co-therapies designed to complement existing immunomodulatory and remyelinating regimens in MS and related neurodegenerative disorders.

## INTRODUCTION

Multiple sclerosis (MS) is a primary idiopathic demyelinating disorder of the central nervous system (CNS), pathologically defined by progressive neurodegeneration, axonal loss, and reactive gliosis (Campbell and Mahad, 2018a; Mahad et al., 2015). As the disease advances into secondary progressive MS (SPMS), acute inflammatory activity wanes and is replaced by chronic microglial activation, oxidative stress, mitochondrial dysfunction, and intracellular ion dysregulation(Campbell and Mahad, 2018b). Together, these compartmentalized mechanisms drive steady myelin degradation and axonal loss, generating the multifocal CNS lesions and cumulative disability characteristic of progressive disease(Gouider et al., 2024; Mahad et al., 2015; Thompson et al., 2018; Waubant et al., 2019).

Morphologically, axonal injury manifests as focal varicosities or “blebs”, which form when metabolic failure halts fast axonal transport, triggering an accumulation of organelle cargo and biological debris(Mahad et al., 2009; Povlishock, 1992; Summers et al., 2014; Trapp et al., 1998; van den Berg et al., 2017). Concurrently, localized inflammatory cascades dismantle key cytoskeletal elements, including microtubules and neurofilament light chain (NfL), the primary structural subunit supporting myelinated axon integrity. When axonal membranes collapse or transection occurs, disassembled NfL fragments diffuse into the interstitial fluid and cerebrospinal fluid (CSF), eventually draining into the systemic circulation. Consequently, elevated serum NfL (sNfL) has emerged alongside retinal nerve fiber layer (RNFL) thinning on optical coherence tomography (OCT), visual evoked potential (VEP) alterations, and Expanded Disability Status Scale (EDSS) progression as a key clinical biomarker for monitoring ongoing neurodegeneration(Chatziralli et al.; Diem et al., 2003; Petzold et al., 2017; Siller et al., 2019) (Kurtzke, 1955).

Preventing irreversible axonal degeneration requires targeting the fundamental molecular cascades driving neuronal loss (Mahad et al., 2015; Neumann et al., 2002; Waxman, 2006). However, therapeutic strategies in MS aimed at upstream interventions, such as suppressing chronic inflammation, promoting remyelination, inhibiting Ca^2+^-activated calpains, or bolstering mitochondrial energy metabolism, have yielded limited clinical success(Cadavid et al., 2019; Chataway et al., 2020; Cree et al., 2020; Trager et al., 2014). These shortcomings highlight the need to add therapies that also target a more distal, convergent node in the axon loss pathway: sterile alpha and TIR motif-containing protein 1 (SARM1) (Essuman et al., 2017; Gerdts et al., 2015; Ko et al., 2020; Murata et al., 2018).

Disruption of axonal transport, whether induced by cytoskeletal degradation, ionic imbalance, or energy failure, depletes the labile axonal survival factor nicotinamide adenylyltransferase 2 (NMNAT2) faster than it can be resupplied via somatic transport(Gilley and Coleman, 2010). The resulting elevation in the nicotinamide mononucleotide to nicotinamide adenine dinucleotide (NMN/NAD^+^) ratio is sensed by the autoinhibitory ARM domain of SARM1, relieving inhibition and driving TIR-domain oligomerization(Essuman et al., 2017; Gerdts et al., 2015; Ko et al., 2020; Murata et al., 2018). Once activated, SARM1 rapidly destroys remaining axonal NAD+, precipitating local metabolic collapse and axon fragmentation, the defining molecular execution points of Wallerian and Wallerian-like degeneration(Coleman and Hoke, 2020; Osterloh et al., 2012).

Because SARM1 functions downstream of diverse pathogenic insults, it offers a single, disease-mechanism-agnostic point of intervention(Krauss et al., 2020). This model is supported by SARM1 knockout studies in experimental autoimmune encephalomyelitis (EAE), which demonstrate preserved axonal structure during early inflammatory phases when transport failure occurs(Liu et al., 2021a; Viar K, 2020). Notably, while NMNAT2 overexpression protects axons in models of retinal degeneration(Tribble et al., 2024) it fails in EAE-associated optic neuritis (Liu et al., 2021b), likely because supplementing an upstream, labile enzyme cannot prevent SARM1 activation once transport failure reaches a critical threshold. Direct SARM1 inhibition bypasses this limitation by blocking the downstream executioner itself, regardless of the initial upstream driver. Small-molecule SARM1 inhibitors have recently been developed and are advancing into clinical trials for peripheral axonopathies (Bratkowski et al., 2022; Feldman et al., 2022; Hughes et al., 2021; Khazma et al., 2022; Leahey et al., 2025), offering a clear translational path for MS. Since axonal degeneration not demyelination itself, is the principal driver of permanent clinical disability, targeting SARM1 represents a compelling neuroprotective strategy to complement existing immunomodulatory treatments.

This therapeutic promise, however, has recently been complicated by reports that a major class of orthosteric SARM1 inhibitors base-exchange inhibitors (BEIs), which act as prodrugs that SARM1 itself converts into active-site-blocking NAD+ analogs can paradoxically *activate* rather than suppress SARM1 when target occupancy is incomplete, accelerating axonal degeneration and neurofilament light chain release both in vitro and in EAE in vivo(Leahey et al., 2025; Mani et al., 2025). This concentration-dependent, bidirectional pharmacology, in which the same compound is protective at saturating occupancy but pro-degenerative at sub-inhibitory doses, underscores that achieving uniform, sustained target engagement rather than simply reducing SARM1 activity on average may be essential for the safe clinical translation of SARM1-directed therapeutics, and highlights the importance of mechanistically characterizing any candidate inhibitor’s dose–response behavior before advancing it toward therapeutic use.

In this study, we evaluated SARM1 pathology across optic nerve crush (ONC) and EAE models using global knockout SARM1-/-, retina-targeted adeno-associated virus (AAV)-mediated gene silencing, and the small-molecule SARM1 inhibitor 5-iodoisoquinoline (5IIQ/DSRM-3715)(Hughes et al., 2021). Global SARM1-/- mice displayed a complex phenotype combining localized neuroprotection with elevated systemic and central cytokine levels, retina-restricted SARM1 knockdown mitigated neurodegeneration without provoking systemic immune alterations. Furthermore, pharmacological inhibition with 5IIQ preserved axonal architecture, reduced serum NfL levels, and restored electrophysiological visual function in both ONC and EAE, despite persistent demyelination. Together, our findings demonstrate that SARM1 inhibition uncouples axonal degeneration from demyelination and gross inflammation, establishing SARM1 as a viable, targetable convergent point for neuroprotective co-therapies in MS and related axonopathies.

## MATERIALS AND METHODS

### Ethics Statement

All animal experiments were conducted in compliance with the ARRIVE guidelines and the National Institutes of Health (NIH) standards for the care and use of laboratory animals. All procedures were approved by the Institutional Animal Care and Use Committee (IACUC) at the University of California, Riverside (Animal Welfare Assurance #123).

### Mice

C57BL/6J and SARM1-/- (B6.129X1-Sarm1tm1Aidi/J, JAX# 018069) were used for these experiments. All mice were bred and housed in an AAALAC-accredited facility and kept on a 12-hour light/dark cycle with unrestricted access to food and water.

### Postmortem Human Samples

Human cerebellar and hippocampal sections were obtained from the NIH NeuroBioBank and Cleveland Clinic. Brain slices were from 8 “normal” (2 females, 6 males) and 9 MS (5 females and 4 males). Sample details are in Supplemental Table 1.

### Mouse primary neuronal cell cultures and treatment

Primary cortical neurons were isolated from postnatal day 0–1 (P0–P1) C57BL/6J mice of both sexes using established protocols(Feri et al., 2025; Sciarretta and Minichiello, 2010; Tiwari-Woodruff et al., 2006). Dissociated neurons were plated on poly-D-lysine-coated coverslips and cultured in Neurobasal medium supplemented with B27 for 14 days *in vitro* (DIV) prior to experimental treatment. At DIV14, cultures were treated with either vehicle (0.1% DMSO) or 1 μM rotenone for 6 h. Following rotenone or vehicle exposure, media were replaced with fresh Neurobasal/B27 medium containing one of three SARM1 inhibitors (purchased from Sigma-Aldrich and synthesized by Katzenellenbogen lab): CZ-48 (CF3), and 5-Iodoisoquinoline (5-IQ, DSRM-3716) at concentrations of 10 uM. Vehicle- and inhibitor-containing media were refreshed every other day for 5 days. At the end of the 5-day treatment period, media were aspirated, and cultures were rinsed once with phosphate-buffered saline (PBS) and followed by ice-cold 10% neutral buffered formalin (Fisher Scientific, Hampton, NH) for an additional 10 min.

### EAE induction and treatment

Active EAE was induced in 8–12-week-old WT C57BL/6J and SARM1*⁻/⁻* (B6.129X1-Sarm1tm1Aidi/J) (SARM1KO) mice using MOG_35-55_ peptide, as previously described(Hasselmann et al., 2017; Karim et al., 2018; Karim et al., 2019; Sekyi et al., 2021). Animals were monitored and scored for clinical disease severity daily, beginning at 7 days post-induction (dpi), using the following scale: 0, unaffected; 1, complete tail limpness; 2, failure to right when rolled onto its back; 3, partial hind-limb paralysis; 4, complete hind-limb paralysis; 5, moribund or death.

At peak disease, WT EAE mice were stratified by clinical score to ensure balanced disease severity across groups, then assigned to receive either vehicle (EAE+V) or 5IIQ (EAE+5IIQ). 5IIQ was dissolved in 0.1% DMSO and further diluted in Miglyol oil for a final injectable formulation, then administered at 25 mg/kg per animal; dosage was calculated based on average group body weight. Mice received daily subcutaneous injections of drug or vehicle from peak disease until the chronic phase of EAE (days 40-45). A separate cohort of mice immunized with complete Freund’s adjuvant (CFA) alone served as controls (Cntrl) for each experiment.

### ONC and treatment

ONC was performed on 6–8-week-old male C57BL/6J mice (n = 8 per group) on day 0. Mice were anesthetized via intraperitoneal injection of ketamine (100 mg/kg) and xylazine (10 mg/kg), yielding approximately 1–1.5 h of surgical anesthesia. The right eye in each mouse was left untouched and served as normal control. A conjunctival incision was made using spring scissors, beginning inferior to the globe and extending temporally around the left eye. The left optic nerve was exposed and crushed 0.5 mm posterior to the globe using cross-action forceps (Dumont #N5; Fine Science Tools, Foster City, CA, USA), applying three successive 5-second crushes. Ophthalmic ointment was applied to the eye immediately following surgery. A separate cohort of mice underwent identical anesthesia and surgical exposure (the left eye only) without nerve crush to serve as sham controls. ONC animals were subjected to anterograde transport assay with a 2 µl intravitreal injection of 1% cholera toxin b-subunit (CTB) conjugated to AlexaFluor488 (Molecular Probes C22841) dissolved in sterile PBS. Intravitreal CTB injection resulted in rapid uptake and filling of retinal nerve fiber layer (RNFL) axon bundles. Eyes and optic nerves were isolated after 7 days post-injection, cut, and imaged for the fluorescent red dye. Groups of ONC mice were also treated with either vehicle (ONC+V) or the SARM1 inhibitor 5IIQ (ONC+5IIQ) via subcutaneous injections at a dose of 25 mg/kg, beginning on day 0 and continuing daily until the experimental endpoint on day 14.

### Optical coherence tomography

OCT imaging was performed on day 14 as previously described (Sekyi et al., 2021). Mice were anesthetized and maintained under ketamine/xylazine anesthesia as described above. Retinal images were acquired using spectral-domain OCT (Envisu R2200 SD-OCT; Leica/Bioptigen, Deerfield, IL) with an 840 nm light source. For each eye, 1,000 A-scans and 100 B-scans were acquired to generate a single OCT volume; three volumes were captured per eye and averaged for analysis. Retinal structure was assessed in the region lateral to the optic nerve head. Retinal layers were automatically segmented using Bioptigen Diver 3.0 software (Leica Microsystems, Deerfield, IL), with segmentation of the retinal cell layers performed to exclude retinal blood vessels from thickness calculations.

### Electroretinograms and visual evoked potentials

ERG and VEP recordings were performed on day 14 using a handheld multi-species electroretinography (i-vivo, Henderson, NV), as previously described (Feri et al., 2025; Sekyi et al., 2021). Briefly, mice were dark-adapted for 5 h and then anesthetized with 2% isoflurane, which was maintained continuously throughout data acquisition. Stainless steel subdermal electrodes (F-Needle Electrode, F-E2; i-vivo, Henderson, NV) were placed at the base of the tail (ground) and bilaterally at the snout (reference). For ERG recordings, silver-embedded thread electrodes (1.5′ Filament; i-vivo) were positioned over the cornea and held in place with mini contact lenses filled with saline to optimize electrical conductivity between the electrode and cornea. For VEP recordings, subdermal electrodes were inserted 2–3 mm lateral to the midline, bilaterally over the visual cortex.

ERG recordings consisted of 5 flashes delivered at 0.5 Hz (one flash every 2 s) at an intensity of 3,000 mcd·s/m², with 30 ms of pre-stimulus baseline, a 300 ms recording window post-flash, and a background luminance of 0 mcd/m². VEP recordings used the same flash intensity, timing, and background conditions but consisted of 71 flashes delivered at 0.5 Hz. For each animal, a minimum of 5 responses were averaged for ERG and 25 responses for VEP. Traces were filtered to remove 60 Hz line noise and low-pass filtered at 150 Hz in MATLAB (MathWorks, Natick, MA); baselines were adjusted to zero at stimulus onset. VEP traces were additionally smoothed using a third-order Savitzky– Golay filter (50-point window). Peak amplitudes and latencies for both ERG and VEP waveforms were identified and quantified in MATLAB. A cohort of naïve, age-matched mice was recorded under identical conditions to establish baseline values.

### Splenocyte Isolation and Cytokine Analysis

Spleens were harvested from anesthetized mice prior to intracardiac perfusion and mechanically dissociated into single-cell suspensions in cold RPMI 1640 supplemented with sodium pyruvate, L-glutamine, and 10% fetal bovine serum (supplemented RPMI). Red blood cells were lysed with ACK lysis buffer (VWR), and the remaining cells were washed, counted, and resuspended in supplemented RPMI. 200,000 splenocytes/well were plated in 12-well plates, stimulated with 25 µg/mL MOG^35–55^ peptide, and culture supernatants were collected after 48 h. Cytokine and chemokine levels in the supernatants were measured by the Cytokine Core (Indianapolis, IN) using Luminex® MagPix (XID #0239) by the Cytokine Core (Indianapolis, IN).

### Blood collection and serum NfL analysis

Blood was collected via cardiac puncture prior to perfusion and allowed to clot at room temperature for 2 h. Samples were then centrifuged at 1,500 × g for 15 min at 4°C, and the resulting serum was collected and stored at -80°C until analysis. Because of the high levels of NfL in EAE samples as observed in pilot experiments, serum samples were diluted as follows: CFA control, 1:50; EAE+V, 1:1000; EAE+5IIQ, 1:1000. NfL concentrations were measured in duplicate using the NF-light™ Neurofilament Light Serum ELISA (UmanDiagnostics, Umeå, Sweden)(Revendova et al., 2022).

### AAV production and intravitreal injection

AAV2 vectors containing the mouse synuclein-gamma (mSncg) promoter (commonly used for RGC-specific expression) driving Cas9, a SARM1-targeting gRNA (for SARM1 knock down, KD) a scrambled/non-targeting gRNA (control) and separate AAV2-mSncg-SARM1-mCherry construct (for SARM1 overexpression, OE) were designed and produced by the Hope Center Viral Vectors Core at Washington University in St. Louis, as previously described (Wang et al., 2020). Viral titers were determined by real-time PCR. AAV-Cas9 (9.5 × 10¹⁰ vector genomes [vg]/mL) and AAV-gRNA (1.9 × 10¹² vg/mL) were combined at a 2:1 ratio (AAV-Cas9:AAV-gRNA) prior to injection. For SARM1 KD, AAV-Cas9 was combined with AAV-SARM1 gRNA; for SARM1 OE, AAV-mScng-SARM1-mCherry was used alone; and control eyes received AAV-Cas9 combined with the scrambled/non-targeting vector. Intravitreal injections were performed 7 days before the 1^st^ MOG EAE injection as previously described(Looser et al., 2018; Nieuwenhuis and Osborne, 2023). Briefly, C57BL/6J mice (n=6/group) were anesthetized with ketamine (100 mg/kg) and xylazine (10 mg/kg). Under a dissecting microscope, a scleral incision was made 2 mm posterior to the superior limbus using a 27-gauge needle. A beveled 34-gauge needle attached to a Hamilton syringe (10 µL Gastight Syringe, Model 84877) was then inserted through the same incision, and 4 µL of the appropriate AAV vector or vector mixture was injected into the vitreous. Post injection care including antibiotic/anti-inflammatory eye drops were used. Efficiency of AAV injections was confirmed by incorporation of HA-tag, mCherry, and GFP in the optic nerve and retinal flat mounts.

### Perfusions and tissue preparation

Mice were deeply anesthetized with isoflurane and intracardially perfused with ice-cold PBS followed by 10% formalin. Retinas were dissected out for retinal flat mount staining. For cross sections of retina, eyes were enucleated prior to being post-fixed in 10% formalin and cryoprotected in 30% sucrose. Eyes and optic nerves of EAE mice were collected and processed for IHC as previously published (Sekyi et al., 2021).

### Immunohistochemistry

Tissue sections were washed with PBS and permeabilized with a 0.3% Triton-X (Electron Microscopy Sciences) and 2% normal goat serum (NGS) (Sigma-Aldrich) solution. Tissues were blocked with 20% NGS prior to being incubated with primary antibodies (Supplementary Table 2) overnight at 4°C. The following day, the sections were washed with PBS and then 1X Tris buffered saline (TBS). The sections were incubated with secondary antibodies and co-stained with 4’,6-Diamidino-2-Phenylindole (DAPI; EMD Millipore) to quantify cell numbers. Sections were washed, mounted, cover slipped for confocal microscopy.

### Microscopy and quantification

Retina and optic nerve sections were imaged using an Olympus BX61 spinning disk confocal microscope equipped with 10x and 40x Super Apochromat objectives (Olympus America Inc., Cypress, CA) connected to a camera (Hamamatsu Orca-R2). Z-stack images were acquired, and projection images were compiled using Slidebook 6 and cellSense software (Intelligent Imaging Innovations Inc, Santa Monica, CA). Immunofluorescence intensity and cell numbers were assessed with NIH ImageJ software (v1. 50i http://rsb.info.nih.gov/ij/) and quantified as previously published (Sekyi et al., 2024; Sekyi et al., 2021). Results from all counts were analyzed in GraphPad Prism for statistical significance.

### Statistics

Statistical power calculations were performed prior to study initiation to determine adequate sample sizes for each experimental cohort (n=8-10 mice/group for EAE and ONC treatment studies; n=5 mice/group for in vivo electrophysiological, optical imaging, and IHC studies, evaluating both eyes per animal). To minimize experimental bias, drug administrations, clinical EAE scoring, tissue quantification, and data processing were conducted by investigators blinded to treatment conditions using coded identifiers. All experimental procedures were independently replicated at least twice. Longitudinal EAE clinical scores were analyzed using a two-way unbalanced analysis of variance (ANOVA) followed by Bonferroni Post Hoc Test multiple comparisons test to evaluate differences across time points between treatment groups(Hasselmann et al., 2017). For histological quantification, two representative brain sections or three representative retinal/optic nerve sections per mouse were analyzed for each region of interest. Endpoint comparisons across histological (IHC), electrophysiological (VEP, ERG), and structural (OCT) datasets were evaluated using One way ANOVA with Bonferroni’s Post Hoc tests. Both eyes were assessed in all in vivo studies. Two-group comparisons: CFA control vs. EAE, Normal vs. ONC, Sham vs. ONC, and Vehicle-vs. 5IIQ-treated ONC (or Sham) eyes were analyzed using unpaired t-tests with Welch’s correction. VEP/ERG recordings were performed in matched pairs on individual days. For human postmortem tissue two representative brain sections/postmortem brains (n=6-8 subjects/group) were analyzed for each region of interest. Normal brain sections were compared to MS brain sections using unpaired t tests with Welch’s correction. Similarly, all experiments with 2 group comparison (WT versus SARM1KO and CFA ctrl versus EAE serum NfL and peripheral cytokines) were performed using unpaired t tests with Welch’s correction. Differences were considered significant at the *p<0.05, **p<0.01, ***p<0.001, and ****p<0.0001 level.

## RESULTS

### Elevated SARM1 expression in postmortem tissue from progressive multiple sclerosis cases

SARM1 expression has been well characterized in the rodent brain, but evidence in human brain tissue remains limited. Based on data from the Human Protein Atlas(Uhlen et al., 2015), SARM1 mRNA is ubiquitously expressed in the human brain, with detectable levels in regions including the cerebellum and hippocampus. To establish SARM1 expression in MS postmortem tissue, we compared cerebellar and hippocampal sections from a cohort of predominantly progressive MS (PMS) cases (n = 9) to normal control sections (n = 8) (Supplementary Table 1) by immunohistochemistry (Figure 1). Group differences were assessed using unpaired t-tests.

**Figure 1:**
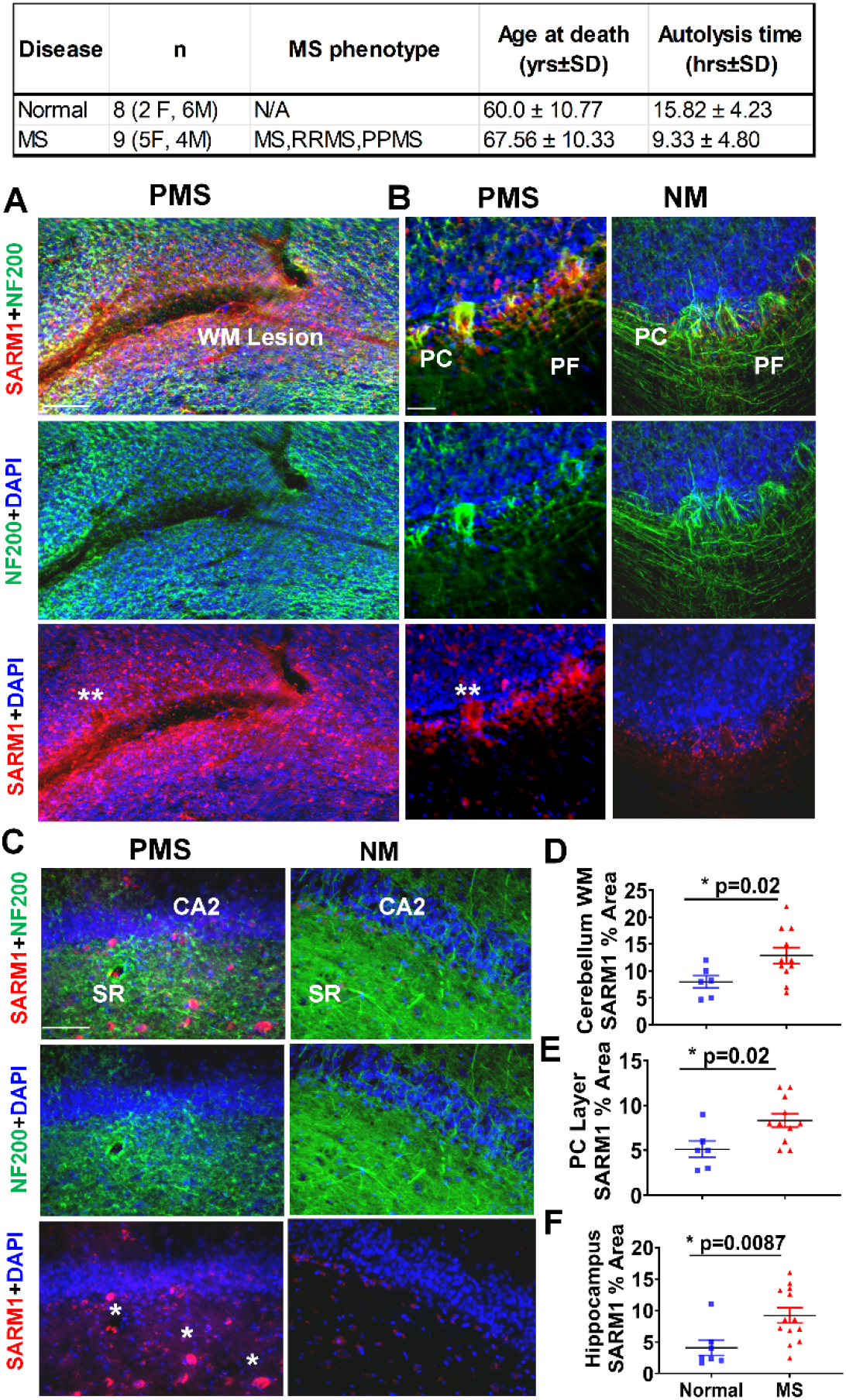
Increased SARM1 immunoreactivity in progressive MS cerebellum and hippocampus. Postmortem cerebellum and hippocampus sections from normal control (NC) and progressive multiple sclerosis (PMS) cases were examined (case details provided in Supplementary Table 1). (**A–B**) Representative confocal images of PMS cerebellum immunostained for SARM1 (red) and NF200 (green). SARM1 expression was increased in white matter (WM; A) and the Purkinje cell (PC) layer (**B**), but not in parallel fibers (PF), alongside a corresponding reduction in NF200 intensity. (**C**) Representative confocal images of PMS and normal control hippocampus, showing elevated SARM1 expression in the CA2 neuronal layer and stratum radiatum (SR) of PMS tissue together with reduced NF200 staining relative to normal control. (**D–F**) Quantification of SARM1 signal intensity showing significant increases in PMS relative to normal control tissue within cerebellar WM (**D**), the PC layer (**E**), and hippocampal CA2/SR (**F**). Data are shown as mean + SEM (blue, normal control; red, PMS). Data represents mean ± SEM (n=8 normal, n=9 MS), Unpaired t test with Welch’s correction, *p<0.05.

PMS cerebellar sections showed significant axonal damage, evidenced by loss of parallel fibers stained with NF200. SARM1 expression was correspondingly increased in both the white matter (WM: t=2.628, df=14.85, p=0.02) and Purkinje cell (PC) layer (t=2.675, df=11.41, p=0.02) relative to controls (Figure 1D–E). Similarly, PMS hippocampal sections showed a significant reduction in NF200 immunostaining compared to normal tissue, while SARM1 expression, which was low in normal hippocampus, was significantly increased near the CA2 region in PMS tissue (t=2.99, df=15.54, p=0.0087) (Figure 1F). Collectively, these findings demonstrate that SARM1 is upregulated in both the cerebellum and hippocampus of postmortem PMS tissue at sites of axonal damage, supporting its regional involvement in disease-associated neurodegeneration.

#### Chronic EAE Produces Optic Nerve Demyelination, Gliosis, and Axon Damage Alongside Elevated Serum NfL and SARM1 Upregulation

Consistent with the axon damage observed in MS(Barton et al., 2019; Costello et al., 2006; Ferguson et al., 1997; Fisher et al., 2006; Garcia-Martin et al., 2010; Green et al., 2010; Herrero et al., 2012; Trapp et al., 1998), comparable damage has been reported in the visual system of the experimental autoimmune encephalomyelitis (EAE) model. Our lab and others have previously demonstrated significant visual pathway pathology in EAE (Jin et al., 2019; Marenna et al., 2020; Mey et al., 2022; Sekyi et al., 2024; Sekyi et al., 2021) with axon damage detectable early in disease course, increasing through mid-disease, and persisting into the late/chronic phase (Sekyi et al., 2024). As proof-of-principle, C576BL/6J mice (n=8) were induced with EAE using the myelin oligodendrocyte glycoprotein (MOG)_35-55_ peptide and at chronic disease, mice were assessed for visual function, histological changes, and molecular markers of axon damage (Figure 2 and Supplementary Figure 1). Clinical signs of disease appeared between days 8–13 post-induction, with peak clinical scores observed between days 18–21 that persisted into the chronic phase (Figure 2Aii), consistent with previously published disease courses(Atkinson et al., 2025; Mangiardi et al., 2011; Sekyi et al., 2021).

**Figure 2:**
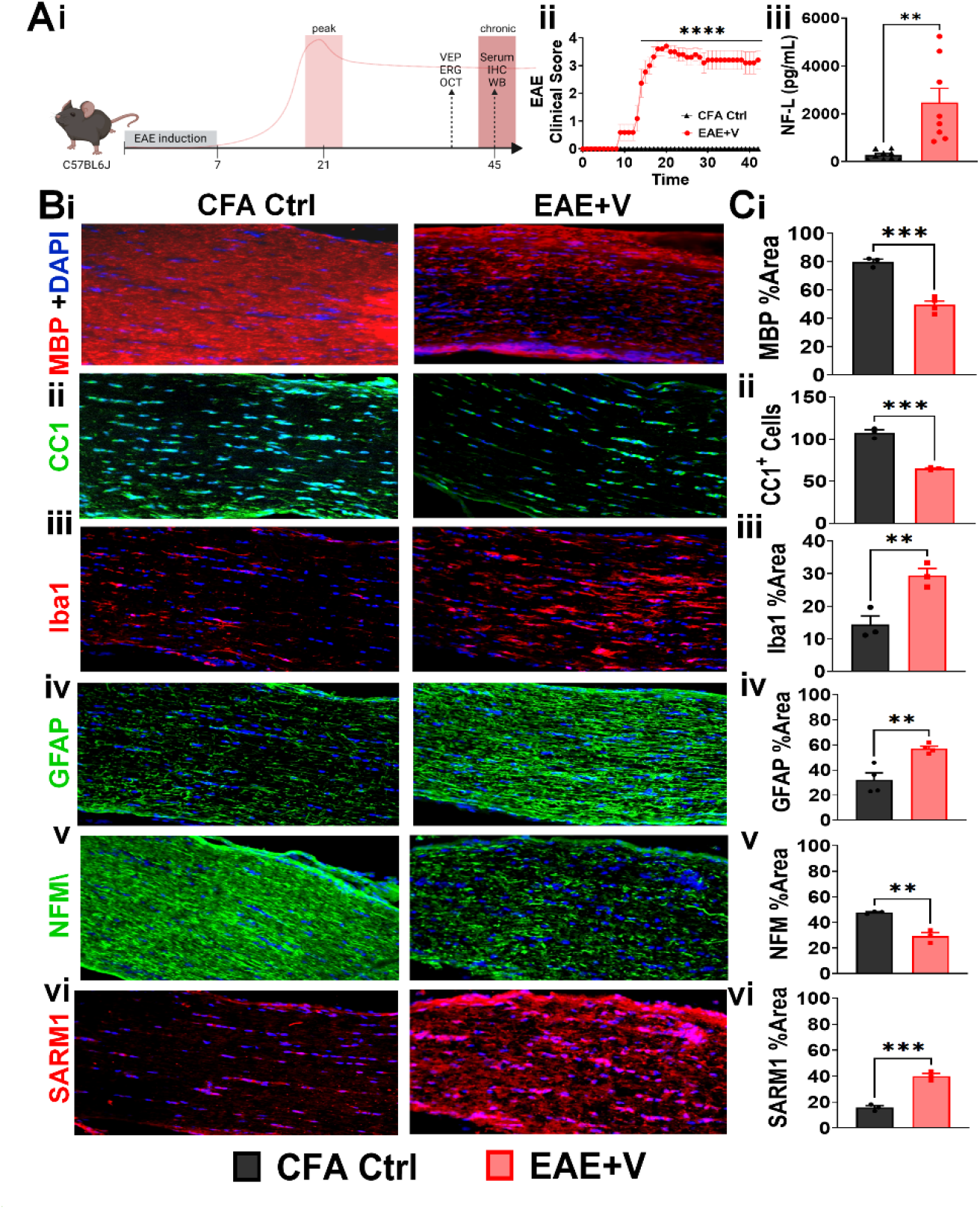
Chronic EAE produces optic nerve demyelination, inflammation, and axon damage alongside elevated serum NfL and optic nerve SARM1 immunoreactivity. **(Ai)** Schematic of EAE induction (MOG35–55) in C57BL/6J mice and the experimental timeline, showing peak disease (∼day 21), functional assessment (VEP, ERG, OCT), and collection of serum, splenocytes for cytokine, and tissue for IHC at chronic disease (∼day 45). **(Aii)** EAE+V mice developed significantly elevated clinical scores relative to CFA controls beginning in the first three weeks post-induction and persisting through the chronic phase. Two-way ANOVA (mixed effect analysis), ****p <0.0001. **(Aiii)** Serum NfL was significantly elevated in EAE+V mice relative to CFA controls at chronic disease. Unpaired t test with Welch’s correction, **p=0.009. **(Bi–vi)** Representative longitudinal optic nerve sections from CFA control and EAE+V mice immunostained for MBP (myelin), CC1 (mature OLs), Iba1 (microglia/macrophages), GFAP (astrocytes), NFM (axonal neurofilament), and SARM1, with DAPI nuclear counterstain. **(Bi–vi)** A significant decrease in MBP **(Bi)** and CC1⁺ cell density **(Bii),** a significant increase in Iba1 **(Biii)** and GFAP **(Biv)**, a significant decrease in NFM **(Cv),** and a significant increase in SARM1 **(Bvi)** immunoreactivity was observed in EAE optic nerve sections as compared to CFA ctrl sections. **(Ci-vi)** Data represents mean ± SEM (black, CFA Ctrl; red, EAE+V; N=6-8 mice/group). Unpaired t test with Welch’s correction **p < 0.01, ***p < 0.001, ****p < 0.0001.

Neurofilament light chain (NfL), a structural neuronal protein, has been widely used as a biomarker of axon damage across neurodegenerative diseases. In both MS and EAE (Ferreira-Atuesta et al., 2021; Galetta et al., 2021) elevated serum or plasma NfL is a promising indicator of axonal damage and disease progression. To determine whether axonal damage could be detected by serum NfL in our model, serum was collected from EAE and control mice at chronic disease and analyzed using the NF-Light Serum ELISA. Serum NfL was significantly elevated in EAE mice relative to CFA controls (t=3.811, df=15, p=0.0017) (Figure 2Aiii).

Our lab has previously reported extensive OCT, ERG, and VEP deficits in EAE mice (Sekyi et al., 2024; Sekyi et al., 2021). Consistent with these findings, OCT imaging revealed significant RNFL thinning in EAE mice compared to CFA controls (t = 4.105, df = 11, p = 0.0009) (Supplementary Figure 1Ai-ii). ERG recordings showed decreased A-wave (t = 2.871, df = 28, p = 0.0039) and B-wave (t = 2.601, df = 29, p = 0.0072) amplitudes, along with increased A-wave (t = 2.347, df = 34, p = 0.0125) and B-wave (t = 1.731, df = 29, p = 0.0471) latencies, in EAE mice relative to controls (Supplementary Figure 2Bi–iii). VEP recordings similarly showed decreased P1 (t = 4.974, df = 30, p < 0.0001) and N2 (t = 6.366, df = 36, p < 0.0001) amplitudes, along with increased P1 (t = 3.792, df = 44, p = 0.0002) and N2 (t = 5.822, df = 31, p < 0.0001) latencies (Supplementary Figure 2Biv–vi). Together, these functional deficits confirm that visual pathway axon damage is present in EAE mice.

Optic nerve pathology in EAE is characterized by demyelination, oligodendrocyte (OL) loss, gliosis, and impaired axonal integrity (Sekyi et al., 2024; Sekyi et al., 2021). During axon damage, neurofilaments are fragmented and individual subunits are released into the extracellular space(Yuan and Nixon, 2021). To determine the histopathological basis of this injury, longitudinal optic nerve sections from CFA control and EAE mice were immunostained for markers of myelination (MBP, CC1), inflammation (Iba1, GFAP), and axonal integrity (NFM, SARM1) (Figure 2B). EAE optic nerves showed significantly reduced MBP and CC1⁺ oligodendrocyte staining relative to CFA controls, indicating substantial demyelination and oligodendrocyte loss (Figure 2Ci–ii). This was accompanied by significantly increased Iba1⁺ microglial/macrophage and GFAP⁺ astrocyte staining, reflecting robust glial activation (Figure 2Ciii–iv). Consistent with axonal injury, NFM staining was significantly reduced in EAE optic nerves compared to CFA controls (Figure 2Cv). Furthermore, while SARM1 immunoreactivity was undetectable in CFA control optic nerves, it was significantly increased in EAE optic nerves (Figure 2Cvi). Together, these findings establish that chronic EAE produces coordinated demyelination, glial activation, and axonal cytoskeletal damage in the optic nerve, occurring alongside a marked increase in SARM1 expression that positions this pathway as a candidate driver of the accompanying axon loss.

#### SARM1 ablation reduces but does not eliminate axon damage at peak disease

To assess the effect of SARM1 ablation on EAE clinical disease progression, and SARM1KO mice (n = 8/group) were induced with EAE using MOG_35-55_ peptide and compared to CFA-injected controls. SARM1KO EAE mice showed an earlier onset of clinical disease compared to WT EAE mice; however, by day 20, clinical scores were comparable between the two genotypes (Figure 3A). Similar results were seen earlier(Viar K, 2020).

**Figure 3:**
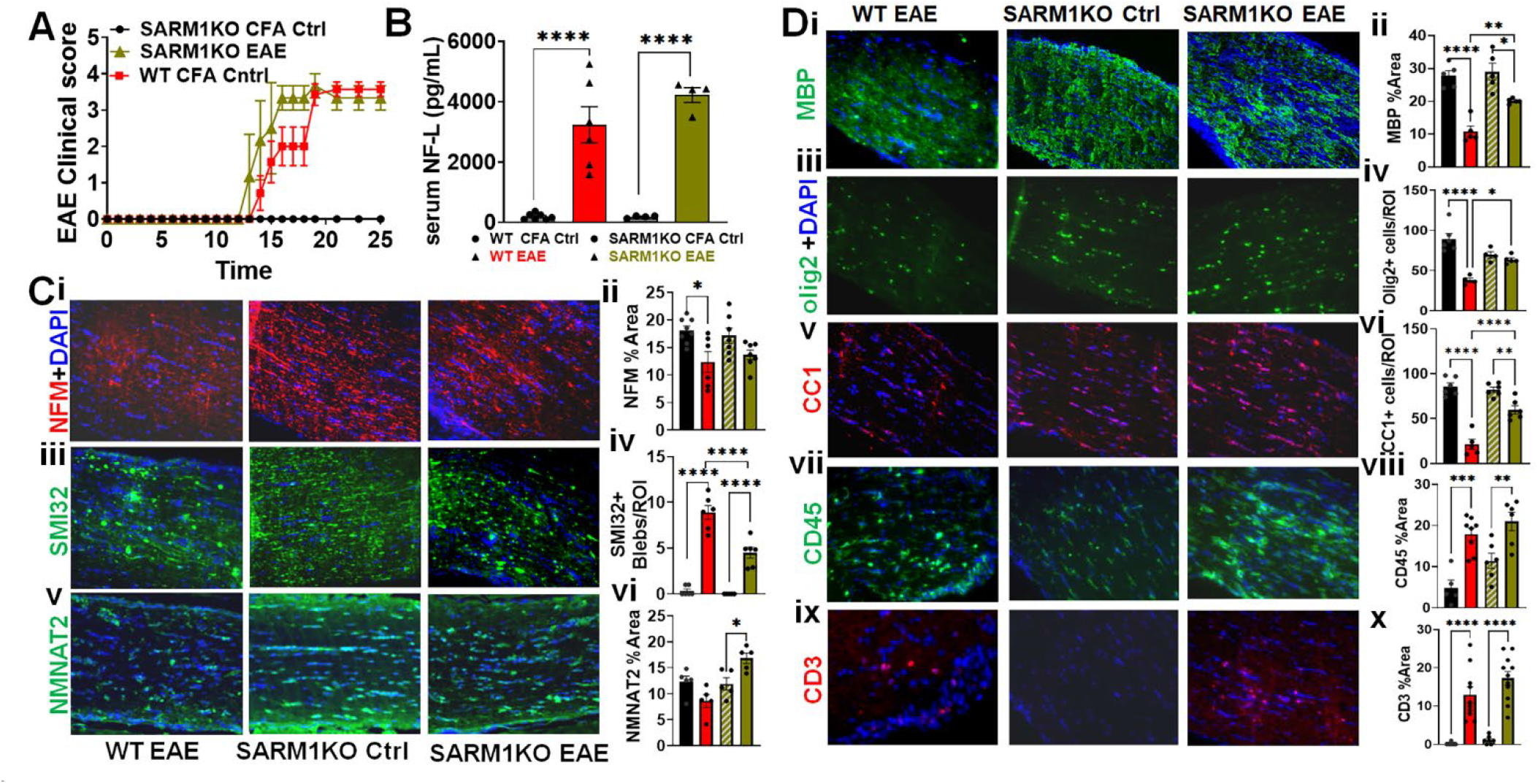
SARM1 knockout (SARM1KO) mice show clinical disease and inflammatory infiltration comparable to WT, but attenuated axon degeneration and reduced OL loss during EAE. **(A**) MOG35–55 EAE was induced in WT and SARM1KO mice. Clinical scores for WT EAE and SARM1KO EAE mice were comparable between genotypes, and both were elevated relative to their respective WT CFA and SARM1KO CFA control groups. **(B**) Serum NfL, assessed on day 25 post-induction, was significantly elevated in both WT EAE and SARM1KO EAE mice relative to their respective CFA controls, with no significant difference between genotypes. Unpaired t test with Welch’s correction, **p=0.0005. **(C)** Representative optic nerve sections immunostained for NFM, SMI-32, and NMNAT2. SARM1KO EAE mice showed no change in NFM compared to SARM1KO CFA controls but showed significantly fewer SMI-32⁺ axonal blebs and increased NMNAT2 expression relative to WT EAE. (**D**) SARM1KO EAE mice showed improved myelination (MBP, Olig2, CC1) compared to WT EAE, without a corresponding change in inflammatory infiltrate (CD45, CD3) (**C-D**) Data represents mean + SEM (N=5–6 mice/group). One-way ANOVA with Bonferroni’s multiple comparisons test, *p<0.05, **p<0.01, ***p<0.001, ****p<0.0001.

Serum NfL was measured after day 25 to assess the extent of axonal damage during EAE progression (Figure 3B). Both WT EAE and SARM1KO EAE mice showed significantly elevated serum NfL compared to their respective CFA controls (WT EAE vs. WT CFA: F(3,11) = 84.22, p < 0.0001; SARM1KO EAE vs. SARM1KO CFA: F(3,11) = 84.22, p < 0.0001). However, serum NfL did not differ significantly between WT EAE and SARM1KO EAE groups, indicating that SARM1 ablation did not reduce this systemic marker of axonal injury.

### SARM1 Ablation Preserves Optic Nerve Cytoskeletal Integrity Despite Elevated Serum NfL

Given that EAE induction produced substantial axonal injury in both WT and SARM1KO mice, as reflected by elevated serum NfL, we next characterized the structural and pathological changes underlying this injury using immunohistochemistry on longitudinal optic nerve sections (Figure 3C). NFM staining, a marker of overall axonal structural integrity, was significantly decreased in WT EAE optic nerves compared to WT CFA controls (F(3,24) = 4.816, p = 0.0173) (Figure 3Ci–ii). NFM staining was comparable between SARM1KO CFA and WT CFA controls. In contrast to WT EAE mice, SARM1KO EAE mice showed no change in NFM staining relative to SARM1KO CFA controls, indicating that SARM1 ablation preserved axonal cytoskeletal structure despite EAE induction.

To further assess axonal stress and injury, we quantified non-phosphorylated neurofilament H (SMI-32), a marker of axonal blebbing and axon damage. Both WT EAE and SARM1KO EAE mice showed a significant increase in SMI-32⁺ axonal blebs compared to their respective CFA controls (F(3,19) = 64.70, p < 0.0001) (Figure 3Ciii–iv). Notably, SARM1KO EAE mice showed significantly fewer SMI-32⁺ blebs than WT EAE mice (F(3,19) = 64.70, p < 0.0001), indicating that SARM1 ablation partially protects against axonal blebbing during EAE, despite not preventing it entirely.

### SARM1 Ablation Is Associated with Increased NMNAT2 Levels During EAE

NMNAT2 lies upstream of SARM1 signaling and plays an essential role in maintaining NAD+ levels and suppressing SARM1 activation(Carty and Bowie, 2019; Gilley and Coleman, 2010). To determine whether SARM1 ablation affects NMNAT2 expression, optic nerve sections were stained for NMNAT2 (Figure 3Cv–vi). NMNAT2 levels were unchanged in WT EAE mice compared to WT CFA controls. In contrast, SARM1KO EAE mice showed significantly increased NMNAT2 levels compared to SARM1KO CFA controls (F(3,16) = 8.671, p = 0.0361). Together, these findings suggest that loss of SARM1 during EAE promotes an increase in NMNAT2, potentially reflecting a compensatory feedback mechanism within the NAD+ homeostasis pathway when downstream SARM1 signaling is absent.

#### SARM1 ablation reduced demyelination but not inflammation at peak disease

EAE evokes an immune response that drives significant demyelination. To determine whether SARM1 ablation affects inflammatory demyelination, longitudinal optic nerve sections were stained for markers of myelination and inflammation (Figure 3D). WT EAE mice showed significant demyelination and OL loss, with decreased myelin basic protein (MBP; F(3,16) = 23.54, p < 0.0001), OL transcription factor 2 (Olig2; F(3,15) = 18.3, p < 0.0001), and anti-adenomatous polyposis coli clone (CC1; F(3,19) = 43.27, p < 0.0001) compared to WT CFA controls (Figure 3Di–vi). In contrast, SARM1KO EAE mice showed a small but significant difference in MBP (F(3,16) = 23.54, p=0290) and CC1 (F(3,19) = 43.27, p =0.0021) but not olig2 compared to SARM1KO CFA controls, indicating that SARM1 ablation prevented EAE-induced demyelination and OL loss. Consistent with this, SARM1KO EAE mice showed significantly higher MBP (F(3,16) = 18.62, p = 0079), Olig2 (F(3,15) = 18.3, p = 0.0144), and CC1 (F(3,19) = 43.27, p < 0.0001) staining compared to WT EAE mice.

Despite this protection against demyelination, SARM1 ablation did not reduce CNS inflammation. WT EAE mice showed significant inflammatory infiltration, with increased CD45 (F(3,21) = 13.53, p = 0.0086) and CD3 (F(3,32) = 27.15, p < 0.0001) staining compared to WT CFA controls (Figure 3Dvii–x). Similarly, SARM1KO EAE mice showed comparable increases in CD45 (F(3,21) = 13.53, p = 0.0004) and CD3 (F(3,32) = 27.15, p < 0.0001) compared to SARM1KO CFA controls, indicating that SARM1 ablation preserves axon and myelin integrity independently of, and without altering, the underlying inflammatory response.

Overall (data summarized in Supplementary Table 3) SARM1 ablation confers substantial but incomplete structural axon and myelin protection during EAE. This occurs independently of any reduction in clinical severity, serum NfL, or CNS/peripheral inflammatory infiltration.

#### SARM1KO Splenocytes Show a Selective, Bidirectional Cytokine and Chemokine Response to EAE

Beyond its role in axon degeneration, SARM1 is known as a negative regulator of inflammatory signaling within the MyD88 family of TLR adaptor proteins (Carty and Bowie, 2019; Sarkar et al., 2023). Given that there was axonal protection, but no significant alteration to CNS immune infiltration in SARM1KO EAE mice compared to controls, we next sought to determine how germline SARM1 knockout alters peripheral immune activity in EAE. To do this, splenocytes from SARM1KO CFA control and SARM1KO EAE mice were isolated and restimulated ex vivo with MOG35–55 peptide, and culture supernatants were analyzed by Luminex® MagPix multiplex cytokine/chemokine assay (Figure 4). SARM1KO EAE splenocytes had significantly increased release of the pro-inflammatory cytokines IFN-γ, TNF-α, IL-3, IL-6, and IL-17, as well as the myeloid growth factor GM-CSF, relative to SARM1KO CFA controls (t=2.784, df=5, p = 0.05 to p<0.001). Among chemokines, MCP-1 (CCL2), MIP-1β (CCL4), and CXCL10 were each significantly elevated in SARM1KO EAE splenocytes, and the anti-inflammatory cytokine IL-10 was also significantly increased. In contrast, a distinct subset of mediators IL-9, RANTES (CCL5), MIP-2 (CXCL2), and CXCL1 were significantly *decreased* in SARM1KO EAE relative to SARM1KO CFA controls, with the most pronounced reduction observed for CXCL1 (p < 0.0001). IL-1β, IL-1α, MIP-1α (CCL3), and VEGF did not differ significantly between groups. Together, these findings demonstrate that EAE induction in SARM1KO mice does not produce a uniform amplification of the peripheral immune response; rather, it drives a selective, bidirectional shift in which the majority of pro-inflammatory cytokines and several chemokines rise with disease, while a specific subset of chemoattractant (IL-9, RANTES, MIP-2, CXCL1) already elevated at the SARM1KO baseline relative to WT ironically declines with EAE induction, consistent with a disease-associated normalization of these mediators toward levels more comparable to those seen in WT EAE.

**Figure 4:**
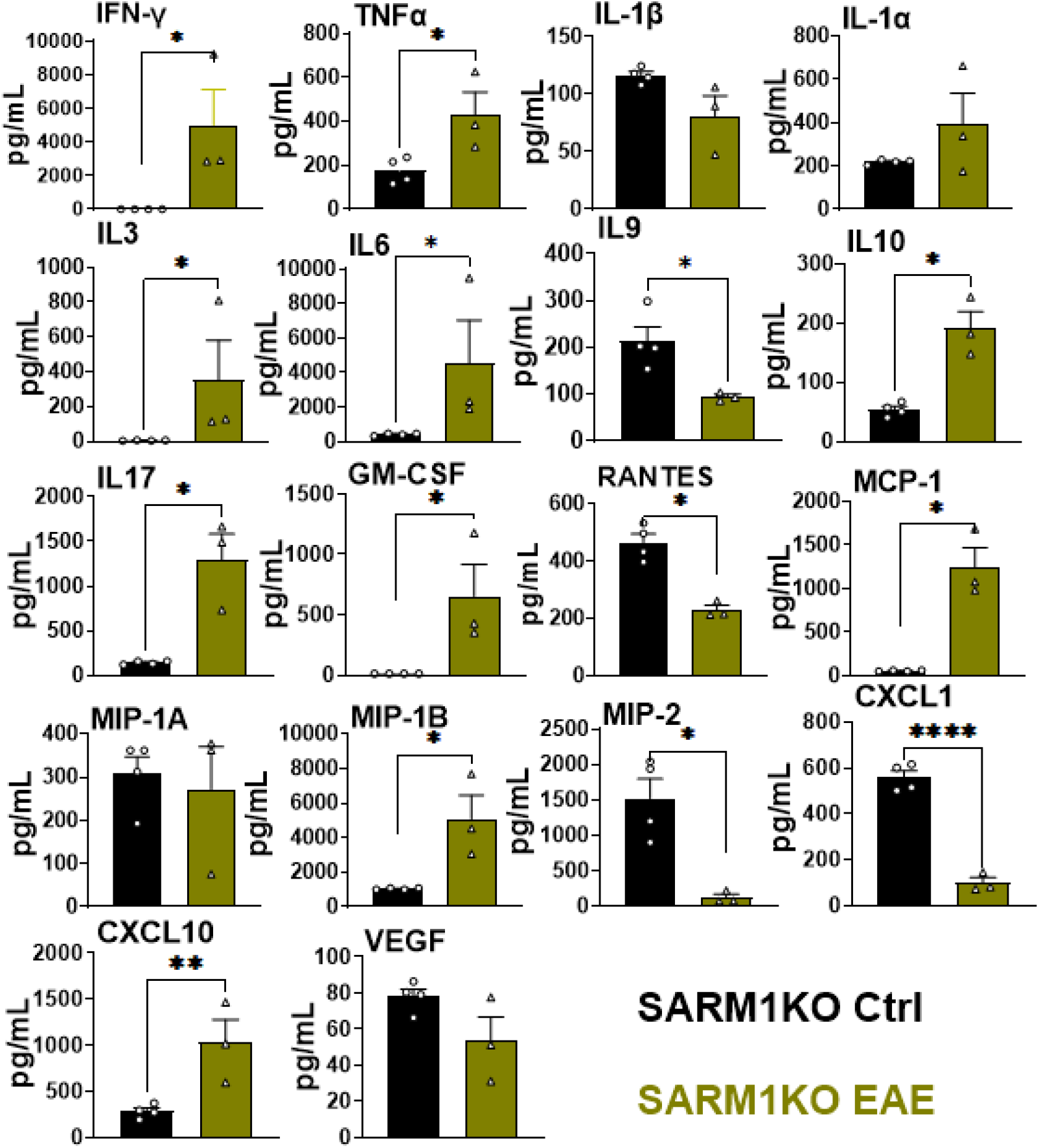
EAE induction in SARM1 knockout (SARM1KO) mice produces a selective, bidirectional shift in peripheral cytokine and chemokine output. Splenocytes from SARM1KO CFA control and SARM1KO EAE mice were isolated and restimulated ex vivo with MOG35–55 peptide, and culture supernatants were analyzed for cytokine and chemokine content. Several pro-inflammatory mediators were significantly increased in SARM1KO EAE relative to SARM1KO CFA controls, including IFN-γ, TNF-α, IL-3, IL-6, IL-17, GM-CSF, MCP-1, MIP-1β, and CXCL10, as well as the anti-inflammatory cytokine IL-10. In contrast, RANTES (CCL5), CXCL1, and MIP-2 (CXCL2) were significantly decreased in SARM1KO EAE relative to SARM1KO CFA controls, and IL-9 was likewise significantly reduced. IL-1β, IL-1α, MIP-1α, and VEGF did not differ significantly between groups. Data represent mean ± SEM (black, SARM1KO Ctrl; olive, SARM1KO EAE) n=3–4 mice/group). Unpaired t test with Welch’s correction, *p<0.05, **p<0.01, ***p<0.001, ****p<0.0001.

#### AAV2-Mediated CRISPR Knockdown and Overexpression of SARM1 in RGCs During EAE

Given that germline SARM1 knockout is not cell-type specific, we next investigated RGC-specific SARM1 manipulation during EAE, as RGCs are severely affected in this model and provide a tractable system for evaluating axon damage. To knock down (KD) SARM1 in RGCs, we used an AAV2 vector expressing Cas9 under the mouse Sncg (mSncg) promoter together with SARM1-targeting gRNAs(Wang et al., 2020) (Figure 5Ai). SARM1 overexpression (OE) was performed using an AAV2 vector driven by the same mSncg promoter. Mice received an intravitreal injection of AAV mediating SARM1 KD or OE in the left eye, while the contralateral eye received a control scrambled gRNA to serve as an internal control. Seven days post-injection, mice were induced with EAE using MOG_35-55_ peptide. Twenty-eight days post-induction, ERGs and VEPs were performed, followed by tissue collection for IHC (Figure 4Aii). Clinical disease severity was comparable across SARM1 KD, SARM1 OE, and EAE control groups (Figure 5Aiii).

**Figure 5:**
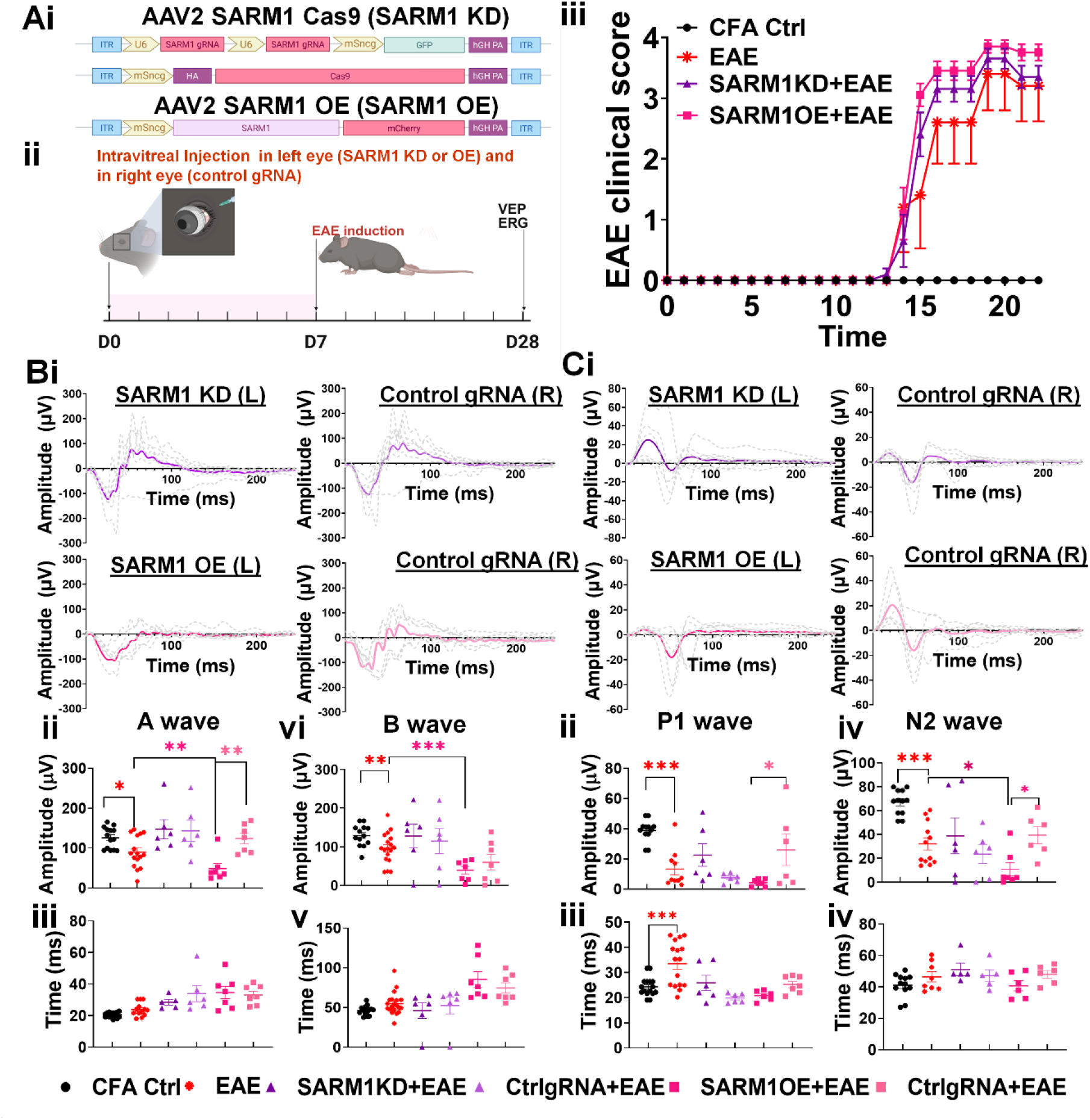
Localized decrease in retinal SARM1 induces decreased functional deficits. (Ai) Schematic of AAV vectors for SARM1 knockdown (KD, AAV2-SARM1-Cas9) or overexpression (OE, AAV2-SARM1OE). (Aii) Timeline: AAV injection, EAE induction 7 days later, visual function assessed 14 days post-induction. (Aiii) Neither SARM1KD nor SARM1OE altered EAE clinical scores relative to EAE controls. (B) ERG traces (individual eyes and group averages) for control (black), EAE (red), SARM1KD (dark purple), and SARM1OE (magenta. SARM1KD showed no change in A-wave amplitude and latency. SARM1OE decreased A-wave amplitude relative to internal gRNA EAE controls. (C) VEP traces showed no changes in SARM1KD induced P1 and N2 amplitude or latency, while SARM1OE reduced P1 amplitudes without changes to N2 amplitude and latencies as compared to internal gRNA control eye. Data represent mean ± SEM (N=5–6 mice/group). One-way ANOVA with Bonferroni’s multiple comparisons test *p<0.05, **p<0.01, ***p<0.001, ****p<0.0001.

To assess how RGC-specific SARM1 KD or OE affects retinal function, ERGs were recorded at peak disease (Figure 5B). Individual traces (dashed lines) and a representative trace (bold) from the AAV2-injected (KD or OE) left eye of each mouse are shown alongside the corresponding internal control (contralateral, control gRNA-injected) right eye (Figure 5Bi). SARM1 KD eyes were associated with increased A-wave and B-wave amplitude in the injected and internal control eyes relative to EAE/gRNA control, indicating that this effect was not specific to local SARM1 knockdown.

In contrast, SARM1 OE produced a change specific to the injected eye: A-wave amplitude was significantly decreased relative to both the internal control eye (F(3,40) = 14.67, p < 0.0001) and the EAE/gRNA control eye (F(3,40) = 14.67, p = 0.0012). Neither SARM1 KD nor SARM1 OE significantly affected A-wave or B-wave latency by this analysis. Unlike SARM1 KD, the OE eye therefore diverged clearly from its internal control, indicating that a local increase in SARM1 substrate produces a functional deficit specific to the manipulated eye, beyond what is seen in the contralateral eye or in EAE/gRNA controls (Figure 5Bii-v).

Together, these findings indicate that RGC-specific SARM1 manipulation during EAE produces asymmetric functional consequences: SARM1 overexpression is sufficient to worsen retinal function in a locally restricted manner, whereas SARM1 knockdown does not produce a locally restricted functional benefit. Given that EAE is a systemic autoimmune disease with bilateral inflammatory and demyelinating insult, we interpret this asymmetry as reflecting the practical limits of a partial, unilateral knockdown: incomplete local SARM1 reduction is likely outpaced by the ongoing systemic disease process, such that it is not sufficient to meaningfully improve retinal function in the injected eye relative to its contralateral counterpart whereas even a partial, local increase in SARM1 substrate is sufficient to produce a detectable, eye-specific deficit.

To determine whether RGC-specific SARM1 KD or OE affected visual function beyond the retina, VEPs were recorded. As with ERGs, individual traces (dashed lines) and a representative trace (bold) from the AAV2-injected left eye of each mouse are shown alongside the corresponding internal control eye (Figure 5Ci). By one-way ANOVA, SARM1 OE produced a small but significant decrease in both P1 (F(5,41) = 8.975, p = 0.0224) and N2 (F(5,41) = 8.975, p = 0.0126) amplitude relative to the EAE/gRNA control eye, with no significant effect on P1 or N2 latency across groups. SARM1 KD did not significantly alter P1 or N2 amplitude or latency relative to EAE/gRNA or internal control eyes (Figure 5Cii-v).

Overall, SARM1 KD produced no significant change in visual function by VEP, consistent with its largely non-specific ERG effects, whereas SARM1 OE produced a modest but significant reduction in VEP P1 and N2 amplitude that parallels its more pronounced effect on ERG A-wave amplitude. Together with the histological findings, these results indicate that RGC-restricted SARM1 overexpression produces a consistent, if graded, pattern of functional impairment across the visual pathway most robust at the level of the retina (ERG) and detectable, though more modest, further downstream (VEP) whereas SARM1 knockdown does not produce a reliable functional benefit at either level despite its partial structural protection.

#### Histopathological outcomes of AAV2-mediated CRISPR Knockdown and Overexpression of SARM1 in RGCs During EAE

Successful intravitreal delivery and targeted expression were confirmed by analyzing optic nerves and validating the respective consequences of SARM1 overexpression (mCherry) and SARM1 knockdown (HA-tagged green fluorescence). Retinal flat mounts across the experimental groups were evaluated to assess changes in retinal ganglion cell (RGC) survival (Figure 6Ai, Bi-iv). Quantification using NeuN immunohistochemistry demonstrated a significant reduction in RGC density across all EAE groups relative to CFA and control cohorts. Notably, eyes subjected to SARM1 knockdown exhibited significantly attenuated RGC loss compared to those with SARM1 overexpression or gRNA EAE control (F(3, 18) = 17.65, p = 0.0403) (Figure 6C).

**Figure 6:**
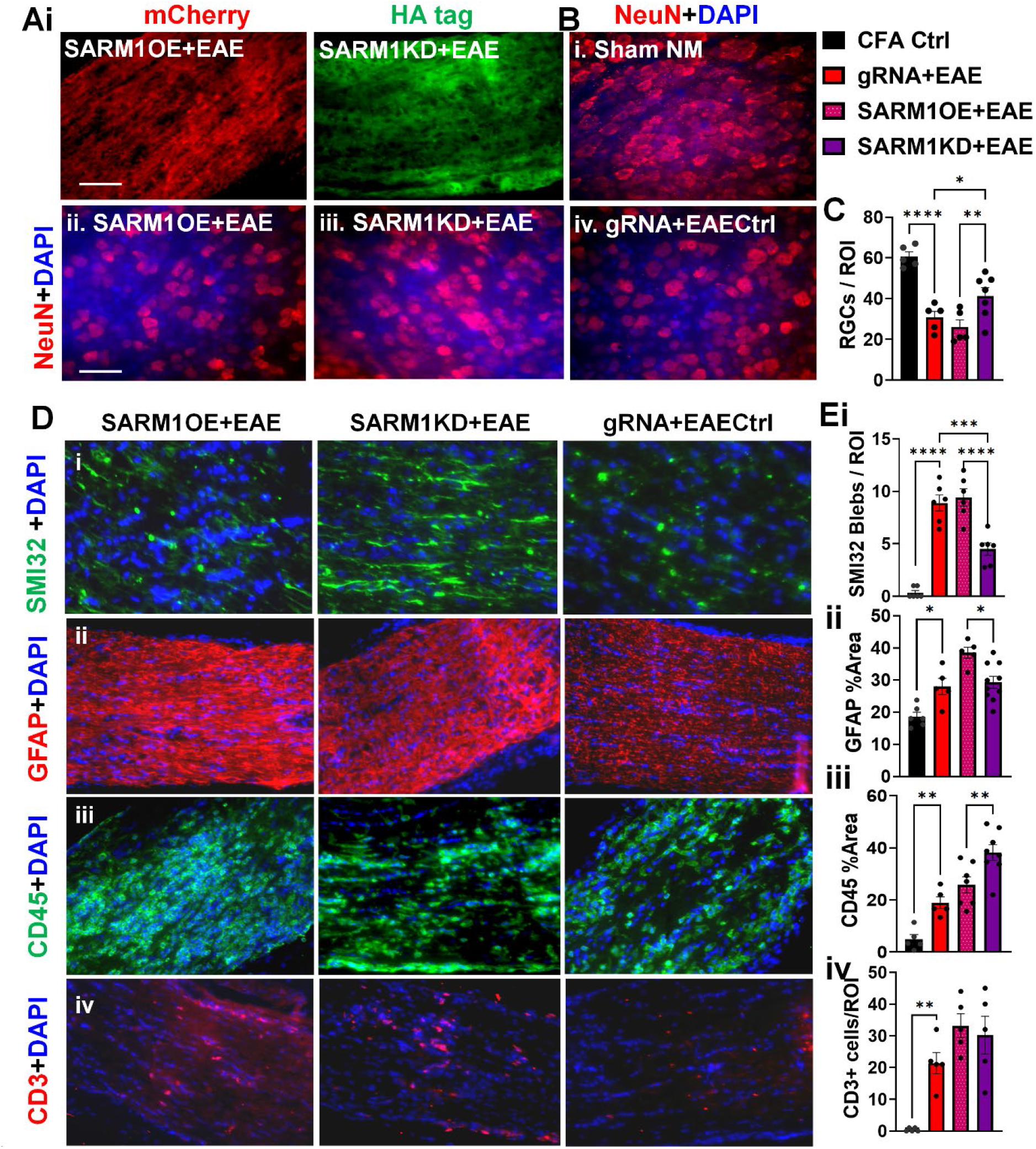
Localized decrease in retinal SARM1 with AAV2 intravitreal injections induces attenuated retinal and optic nerve pathology. Optic nerves and retinas were assessed for AAV2 expression and subsequent consequences of SARM1 manipulation. mCherry (SARM1OE) and HA-tag (SARM1KD) confirmed viral expression in the optic nerve. (E, F) EAE significantly reduced NeuN+ RGCs, an effect reversed by SARM1KD. (G, H) EAE significantly increased SMI32, GFAP, CD45, and CD3 staining; SARM1OE further increased GFAP, while SARM1KD reduced SMI32 and GFAP but increased CD45. Data represent mean ± SEM (N=5– 6 mice/group). One-way ANOVA with Bonferroni’s multiple comparisons test *p<0.05, **p<0.01,***p<0.001, ****p<0.0001.

Axon health was evaluated by SMI-32 immunostaining of optic nerve sections across AAV2 treatment groups. SMI-32⁺ axon blebbing was significantly increased in EAE relative to naïve controls (F(3, 20) = 43. 24, p<0.0001). SARM1 OE optic nerves showed an increase in SMI-32⁺ axon blebs comparable to that seen in EAE control groups, whereas SMI-32⁺ blebbing was significantly lower in the SARM1 KD group (F(3, 20) = 43.24, p = 0.0001) (Figure 6Di, Ei). Notably, the SARM1 KD group also showed numerous elongated, intact SMI-32⁺ axons, consistent with partial preservation of axonal structure.

Inflammation was assessed by quantifying GFAP⁺ astrocytes, CD45⁺ microglia/macrophages, and CD3⁺ T cells in optic nerve sections. All EAE groups showed a significant increase in these inflammatory markers relative to sham CFA controls. Compared to the SARM1 OE group, the SARM1 KD group showed significantly lower GFAP intensity (F(3, 22) = 14.81, p = 0.0130) but significantly higher CD45 intensity (F(3, 21) = 22.83, p = 0.0034) (Figure 6Dii–iii, Eii-iii). CD3⁺ cell numbers did not differ significantly between WT EAE and the SARM1 OE and SARM1 KD groups (Figure 6Div-Eiv).

Overall, SARM1 KD partially preserved RGC number and axonal integrity and selectively reduced astrogliosis relative to SARM1 OE, despite showing higher CD45⁺ myeloid cell presence and no change in CD3⁺ T cell infiltration. In contrast, SARM1 OE was associated with RGC loss and axonal blebbing comparable to EAE controls. Together, these findings (summarized in Supplementary Table 4) indicate that RGC-specific SARM1 knockdown confers partial histopathological neuroprotection that is dissociated from myeloid and T-cell infiltration, whereas SARM1 overexpression provides no protection against EAE-associated axonal and neuronal injury.

### SARM1 Inhibitors Protect Primary Cortical Neurons Against Rotenone-Induced Axonal and Neuronal Damage

Having established the functional and histopathological importance of SARM1 in EAE-associated neurodegeneration, we next asked whether pharmacological SARM1 inhibition could protect neurons against metabolic stress in vitro. Several classes of SARM1 inhibitors are currently under investigation as potential therapies for ALS and other peripheral neuropathies; we selected two readily available compounds, 5IIQ and CF3, to test in primary mouse cortical neurons subjected to mitochondrial stress. Rotenone, an inhibitor of mitochondrial Complex I (NADH-ubiquinone oxidoreductase), was used to model metabolic failure: Complex I inhibition reduces NAD+ levels and increases reactive oxygen species, conditions expected to promote SARM1 activation and axonal degeneration; SARM1 inhibitors were therefore predicted to preserve NAD+ availability and axonal integrity under these conditions.

At 14 days in vitro, primary cortical neuron cultures were treated with vehicle (DMSO) or 1 μM Rotenone for 6 hours, followed by treatment for 5 days with fresh media alone, 10 μM 5IIQ, 10 μM CF3 (Supplementary Figure 2). Wells with no rotenone exposure followed by 10 μM 5IIQ alone, or 10 μM CF3 alone were included as controls. Cultures were then fixed and immunostained for MAP2 (dendritic and somatic marker) and neurofilament (NF200; predominantly axonal marker). Rotenone treatment alone markedly reduced neuronal survival, with extensive loss of MAP2 and NF200 staining (F (5, 55) = 17.04, p<0.0001). Treatment with 5IIQ or CF3 (Compound 4 in (Bosanac et al., 2021)) alone did not significantly alter neuronal morphology or survival compared to vehicle. In contrast, co-treatment with either 5IIQ (F (5, 55) = 17.04, p=0.0002) or CF3 (F (5, 55) = 17.04, p=0.0002) significantly preserved neuronal integrity in the presence of rotenone, with significantly more intact neurons (nearly 50% more) and reduced discontinuous/fragmented MAP2 and NF200 staining compared to rotenone alone; (Supplementary Figure 2). Together, these results demonstrate that pharmacological SARM1 inhibition protects against rotenone-induced axonal damage and neuronal death in vitro.

#### Significant RGC loss, axon damage, and visual dysfunction are evident in ONC mice

To directly assess the effect of SARM1 inhibitors on axon damage, we turned to the ONC model (Supplementary Figure 3). ONC is a well-established model of traumatic optic nerve injury that provides a well-defined system for studying axon damage, as partial or complete disruption of the nerve severs the connection between RGCs and their central targets. ONC produces characteristic axon swellings that can be visualized by intravitreal injection of a tracer such as cholera toxin subunit B (CTB)(Abbott et al., 2013); impaired axonal transport causes biological material, including the tracer, to accumulate within the axon, manifesting as focal swelling or blebbing.

To determine the extent of axon damage in this model, C57BL/6J mice were injected with CTB two weeks prior to ONC injury (Supplementary Figure 3A). ONC was performed in 1 eye, and the other eye was used as a sham control. During 14 days of post-injury (14 dpi), mice were assessed for visual function and ocular histology. ONC mice showed numerous axon swellings/blebs by CTB immunofluorescence compared to sham controls (Supplementary Figure 3B). Retinal flat mounts from different groups were immunostained with NeuN. NeuN⁺ cell counts were significantly reduced in ONC+vehicle (ONC+V) mice compared to sham controls (t = 23.00, df = 14, p < 0.0001) (Supplementary Figure 3C–D), reflecting RGC loss. Consistent with this loss, OCT imaging showed significant RNFL thinning in ONC mice compared to sham controls (t = 2.923, df = 52, p = 0.0026) (Supplementary Figure 3E), in agreement with prior reports linking RGC loss to decreased RNFL thickness.

Concurrent with RGC death and axonal damage, in vivo functional recordings showed significant impairment in visual function, consistent with our prior use of ERG and VEP to assess visual pathway function(Sekyi et al., 2024; Sekyi et al., 2021). ONC significantly reduced ERG A-wave (t = 2.839, df = 7, p = 0.0125) and B-wave (t = 1.910, df = 7, p = 0.0489) amplitudes compared to sham controls (Supplementary Figure 3F). VEP recordings in ONC+V mice showed reduced P1 (t = 3.033, df = 7, p = 0.0095) and N2 (t = 2.558, df = 7, p = 0.0188) amplitudes with no change in latencies.

Taken together, these findings demonstrate that the ONC model recapitulates key features of axon damage pathophysiology, confirming its utility for studying axon damage signaling and for evaluating novel neuroprotective compounds, including SARM1 inhibitors.

#### 5IIQ Treatment Reduces Neuronal Loss in the Retina of ONC Mice

SARM1 ablation provides protection against axon degeneration in various diseases(Geisler et al., 2016; Gerdts et al., 2015; Gerdts et al., 2013; Gilley et al., 2017; Henninger et al., 2016; Marion et al., 2019; Osterloh et al., 2012; Ozaki et al., 2020; Sasaki et al., 2020; Turkiew et al., 2017; Viar K, 2020; White et al., 2019). To determine whether pharmacological inhibition of SARM1 using 5IIQ can protect against axon damage, ONC was performed in one eye, with the contralateral eye serving as a sham control. Immediately following ONC, mice were treated with either vehicle (ONC+V) or 5IIQ at 25 mg/kg/day (ONC+5IIQ) until day 14 (Supplementary Figure 3 and Figure7). Mice were then assessed for functional outcomes, followed by IHC of retinal flat mounts and optic nerve tissue (Figure 7A).

**Figure 7:**
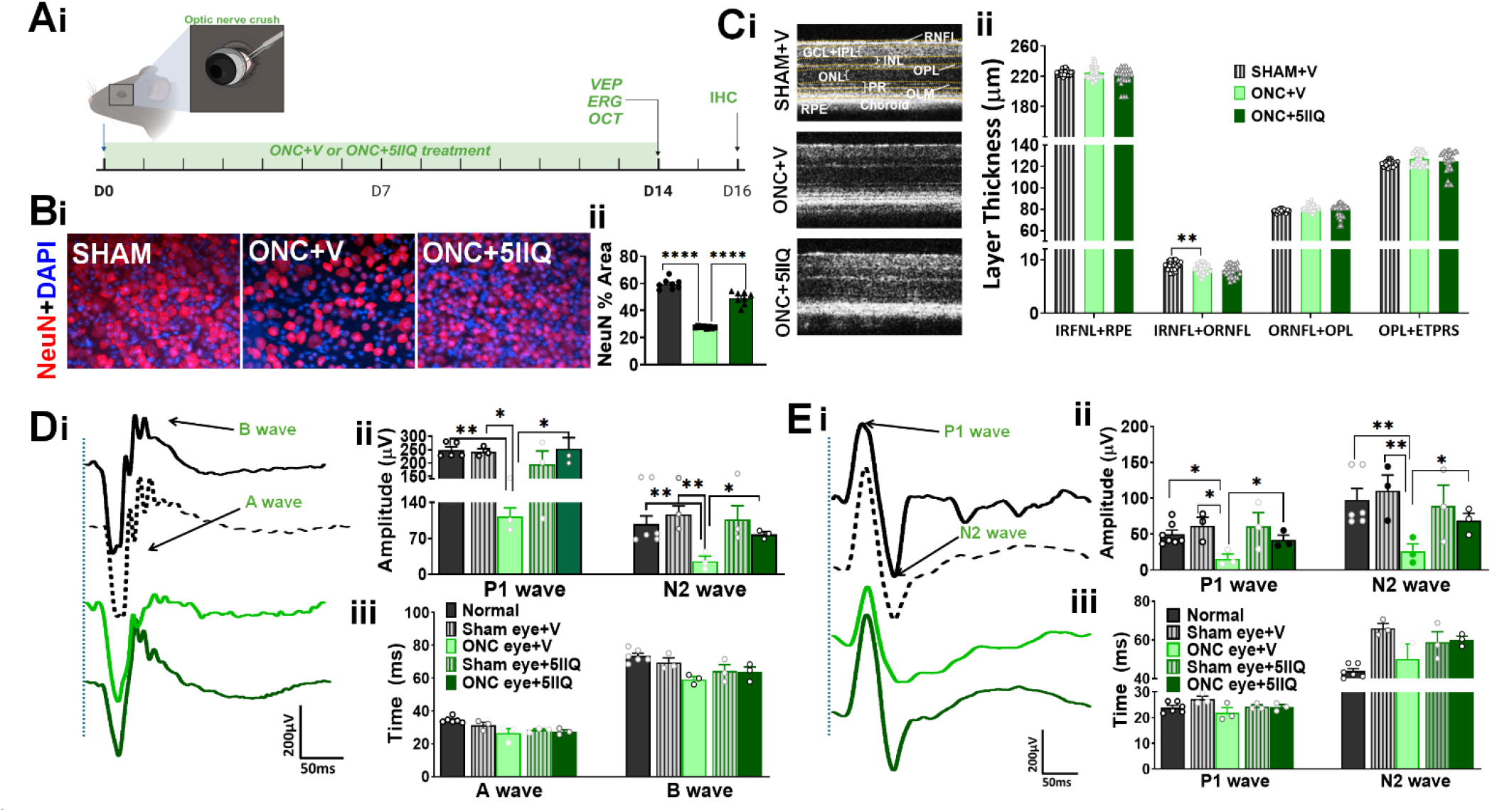
Effect of SARM1 inhibitor, 5IIQ in ONC injury shows significant effects on neuroprotection. (Ai) Schematic of the ONC experimental design. (Bi, ii) Retinal flat mounts show ONC+V significantly decreases NeuN+ RGCs relative to sham, an effect rescued by 5IIQ treatment. (Ci-ii) Representative OCT images across groups. Mice with ONC treated with vehicle have significantly thinner RNFL relative to sham, with no effect of 5IIQ. (Di–iii) Representative ERG traces; ONC does not alter latency but significantly reduces amplitude relative to control, an effect rescued by 5IIQ treatment. (Ei–iii) Representative VEP traces; ONC impairs P1 latency (unaffected by 5IIQ treatment) and significantly reduces amplitude, which 5IIQ treatment rescues. F) Optic nerve sections stained for MBP, Iba1, and GFAP show no effect of ONC without or with 5IIQ treatment on myelination or gliosis. (G) Optic nerve sections stained for NFM, SARM1, and NMNAT2: show significantly reduced NFM and NMNAT2 and increased SARM1 during ONC relative to control. Treatment with 5IIQ shows significant increases in NFM and NMNAT2 but not SARM1. Data represents mean ± SEM (n=6-8 mice/group). One-way ANOVA with Bonferroni’s multiple comparisons test and Unpaired t test with Welch’s correction, *p<0.05, **p<0.01, ***p<0.001, ****p<0.0001.

To assess the extent of RGC survival in ONC mice treated with vehicle or 5IIQ, retinal flat mounts were stained for neuronal nuclear antigen (NeuN), a neuronal marker, 14 days post-injury (Figure 6Bi-ii). Retinas from ONC+V mice showed a significant decrease in NeuN⁺ cells compared to sham controls (F (2, 21) = 167.2, p <0.0001). In contrast, retinas from ONC+5IIQ eyes showed a significant increase in NeuN⁺ cells compared to ONC+V eyes (F (2, 21) = 167.2, p < 0.0001), indicating that 5IIQ treatment preserves RGC survival following ONC (Figure 7Bi-ii).

Consistent with this loss, OCT imaging showed significant RNFL thinning in ONC+V mice compared to sham controls (F (2, 80) = 10.20, p = 0.0001) (Supplementary Figure 3E; Figure 7Cii). Representative OCT images from sham, ONC+V, and ONC+5IIQ mice are shown in Figure 7C. Treatment with 5IIQ did not significantly alter RNFL thickness compared to vehicle-treated ONC mice, indicating that 5IIQ improved retinal and visual pathway function without preventing RNFL thinning.

#### Treatment with 5IIQ Protects Against VEP and ERG Deficits but Not RNFL Thinning in ONC Mice

To determine whether 5IIQ treatment improved visual function in ONC mice, OCT, ERG, and VEP were performed 14 days post-injury. Naïve mice were recorded to establish baseline. Sham controls+V, and ONC+V groups or Sham controls+5IIQ and ONC+5IIQ groups were simultaneously recorded and thus were compared using unpaired t-tests with Welch’s correction. Naïve mice showed no difference in ERG or VEP function compared to sham controls (Figure 7D-E). Sham mice displayed robust ERG responses, with average A-wave and B-wave latencies of 29.87 ms and 66.90 ms, respectively, and average A-wave and B-wave amplitudes of 219.29 μV and 133.98 μV, respectively (Figure 7Di-iii). ONC+V mice showed decreased A-wave (t = 2.839, df = 7, p = 0.0125) and B-wave (t = 1.910, df = 7, p = 0.0489) amplitudes compared to sham mice. Treatment with 5IIQ significantly increased A-wave (t = 3.213, df = 4, p = 0.0163) and B-wave (t = 3.273, df = 4, p = 0.0153) amplitudes in ONC mice compared to vehicle-treated ONC mice (Figure 7Dii–iii).

VEPs were recorded at 14 dpi using previously published protocols (Sekyi et al., 2024; Sekyi et al., 2021). Sham mice exhibited VEP responses with average P1 and N2 latencies of 25.75 ms and 62.47 ms, respectively, and average P1 and N2 amplitudes of 61.81 μV and 110.15 μV, respectively (Figure 7Ei). ONC+V mice showed no changes in latency but a significant decrease in P1 amplitudes (t = 3.033, df = 7, p = 0.0095) and N2 amplitudes (t = 2.558, df = 7, p = 0.0188). Treatment with 5IIQ significantly increased P1 (t = 2.842, df = 4, p = 0.0234) and N2 (t = 2.882, df = 4, p = 0.0225) amplitudes compared to vehicle-treated ONC mice (Figure 7Eii–iii).

### 5IIQ Treatment Protects Against NFM Loss and Increases NMNAT2 in ONC Optic Nerves

In EAE, extensive demyelination and gliosis are observed in the optic nerve(Feri et al., 2025; Sekyi et al., 2024; Sekyi et al., 2021). To determine whether the ONC model, treated with vehicle or 5IIQ, similarly involves demyelination or gliosis, longitudinally sectioned optic nerves were assessed by IHC (Figure 7F). Optic nerves from each group were stained for MBP to assess myelin intensity (Figure 7Fi, iv). Sham control optic nerves displayed robust MBP staining, and no difference in MBP staining was observed in vehicle-or 5IIQ-treated ONC mice compared to sham controls. To assess microglial/macrophage and astrocyte activity, optic nerves were also stained for Iba1 (Figure 7Fii) and GFAP (Figure 7Fiii). Sham optic nerves displayed minimal Iba1 and GFAP immunoreactivity, and neither vehicle nor 5IIQ treatment altered Iba1 or GFAP immunoreactivity compared to sham controls. Together, these data indicate that demyelination and gliosis are not prominent features of the ONC model, and that 5IIQ treatment has no effect on these outcomes.

To determine whether 5IIQ treatment protects axonal cytoskeletal integrity, optic nerves from each group were stained for NFM (Figure 7Gi, iv). Sham optic nerves showed robust NFM staining. NFM staining significantly decreased in vehicle-treated ONC optic nerves compared to sham controls (F (2, 21) = 11.74, p = 0.0005), while 5IIQ treatment significantly improved NFM staining compared to vehicle-treated ONC optic nerves (F (2, 21) = 11.74, p = 0.0015).

A decrease in NMNAT2 following injury leads to activation of SARM1’s NADase activity, depleting NAD+ and causing axon damage. Given the central role of SARM1 and NMNAT2 in axon degeneration signaling, we next assessed how their expression changes with or without 5IIQ treatment. ONC optic nerves treated with vehicle or 5IIQ were stained for SARM1 (Figure 7Gii, v) and NMNAT2 (Figure 7Giii, vi). Sham controls showed minimal SARM1 staining, whereas vehicle-treated ONC optic nerves showed a significant increase in SARM1 expression compared to sham controls (F (2, 21) = 30.04, p < 0.0001). SARM1 expression in 5IIQ-treated ONC optic nerves did not differ significantly from vehicle-treated ONC optic nerves. In contrast, robust NMNAT2 expression was observed in sham controls, and NMNAT2 expression was significantly decreased in vehicle-treated ONC optic nerves compared to sham controls (F (2, 21) = 37.71, p < 0.0001). Treatment with 5IIQ significantly increased NMNAT2 expression compared to vehicle-treated ONC optic nerves (F (2, 21) = 37.71, p < 0.0001).

Together, these data (summarized in Supplementary Table 5) indicate that ONC causes significant axon damage, reflected by decreased NFM staining, which may be driven by the concurrent decrease in NMNAT2 and increase in SARM1 expression. 5IIQ treatment protects against ONC-induced axon damage; notably, this protection is associated with restored NMNAT2 levels, but not with a change in SARM1 expression itself. This is consistent with 5IIQ acting as a functional inhibitor of SARM1 NADase activity rather than a suppressor of SARM1 expression

#### 5IIQ treatment does not alleviate clinical disease severity in EAE mice but improves axon damage

To evaluate the effect of 5IIQ treatment in EAE, C57BL/6J mice were induced with active EAE (Figure 8A) and assessed for visual function, peripheral immune response, serum NfL levels, and retinal and optic nerve pathology following treatment. As described in Figure 2B and published reports, mice displayed initial clinical signs of disease between days 8-13 post-induction, with peak clinical scores occurring between days 18-21. At peak disease, mice were divided into treatment groups and received daily vehicle or 5IIQ (25 mg/kg/day) until chronic EAE (day 48). Clinical disease severity did not differ significantly between 5IIQ- and vehicle-treated mice (Figure 8B).

**Figure 8:**
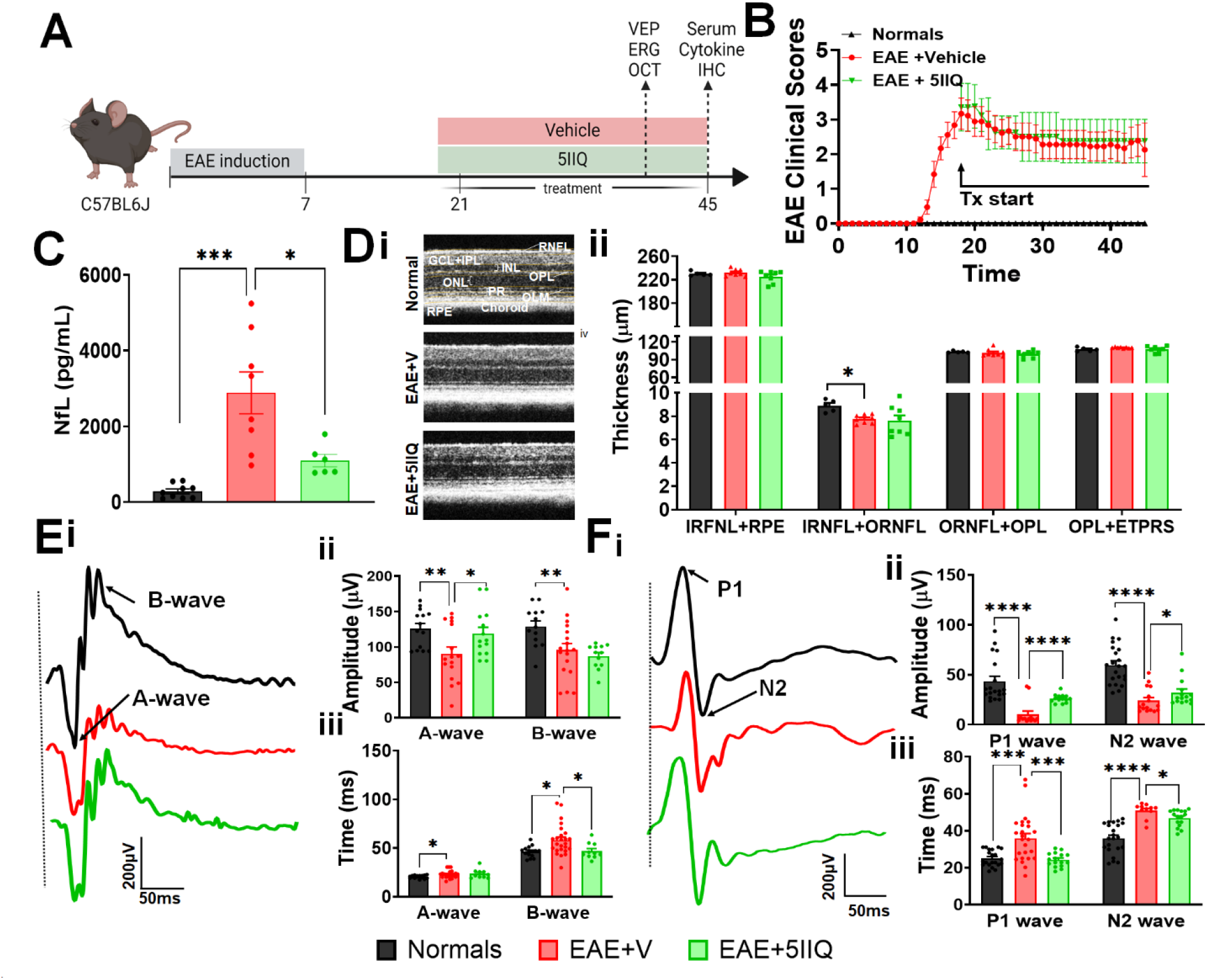
Effect of SARM1 inhibitor, 5IIQ in chronic EAE does not show a decrease in EAE clinical disease but shows a significant effect on axon sparing. (A) Schematic of experimental design. (B) Mice induced with EAE developed clinical disease around Day 8–13, peaking at Day 18–21 when 5IIQ treatment was initiated; 5IIQ treatment did not modify clinical disease, remaining similar to the vehicle-treated group. (C) Serum NfL was significantly elevated in vehicle-treated EAE mice relative to normal controls while treatment with 5IIQ significantly decreased serum NfL levels. (D) OCT imaging shows EAE+V significantly reduced RNFL thickness relative to normal controls. (E) ERG traces show EAE impairs latency and reduced amplitude, while 5IIQ treatment showed significant reversal. (F) VEP traces show EAE impairs latency and reduces amplitude, both of which are improved by 5IIQ treatment. Data represent mean ± SEM (N=5–8 mice/group). One-way ANOVA with Bonferroni’s multiple comparisons test *p<0.05, **p<0.01, ***p<0.001, ****p<0.0001.

To assess the extent of axon damage in treated and untreated animals, serum NfL levels were measured using the NF-Light Serum ELISA assay (Figure 8C). Serum NfL averaged 177.3 pg/mL in normal healthy mice. EAE induction significantly increased serum NfL to an average of 2,353 pg/mL compared to normal mice (F (7, 35) = 5.448, p < 0.0003). Treatment with 5IIQ significantly reduced serum NfL to an average of 1,393 pg/mL compared to vehicle-treated EAE mice (F (7, 35) = 5.448, p = 0.0433).

#### SARM1 Inhibition Does Not Protect Against RNFL Thinning but Alleviates EAE-Induced Functional Deficits

Mice induced with EAE exhibit RNFL thinning and deficits in ERG/VEP responses(Marenna et al., 2020; Mey et al., 2022; Sekyi et al., 2024; Sekyi et al., 2021). To determine how 5IIQ treatment affects retinal structure and function, mice were assessed by OCT and ERG. Representative OCT images from normal, EAE+V, and EAE+5IIQ mice are shown in Figure 7Di. EAE+V mice showed significant RNFL thinning compared to normal mice (F (2, 18) = 3.879, p = 0.0397). RNFL thickness did not differ significantly between 5IIQ- and vehicle-treated EAE mice (Figure 8Dii).

To assess how 5IIQ treatment alters retinal function, ERGs were recorded at chronic EAE (Figure 8Ei–ii). Normal mice showed average A-wave and B-wave amplitudes of 126.1 μV and 129.1 μV, respectively, and average A-wave and B-wave latencies of 46.31 ms and 20.55 ms, respectively. EAE+V mice showed significantly decreased A-wave (F(2,39) = 7.380, p = 0.0013) and B-wave (F(2,39) = 7.380, p = 0.0019) amplitude, along with significantly increased A-wave (F(2,43) = 3.748, p = 0.003) and B-wave (F(2,43) = 3.748, p = 0.005) latency, compared to CFA control mice. Treatment with 5IIQ significantly increased A-wave amplitude compared to vehicle-treated EAE mice (F(2,40) = 6.536, p = 0.0095), with no significant difference in B-wave amplitude. Treatment with 5IIQ significantly decreased A-wave latency (F(2,41) = 6.329, p = 0.0352) and B-wave latency (F(2,41) = 6.329, p = 0.0068) compared to vehicle-treated EAE mice.

To determine how visual function is modified by 5IIQ treatment, VEPs were recorded at chronic EAE (Figure 8Fi–ii). Normal mice displayed robust VEP responses, with average P1 and N2 amplitudes of 43.55 μV and 59.67 μV, respectively, and average P1 and N2 latencies of 25.03 ms and 35.90 ms, respectively. EAE+V mice showed significantly decreased P1 (F(2,47) = 24.17, p < 0.0001) and N2 (F(2,47) = 24.17, p < 0.001) amplitudes, along with significantly increased P1 (F(2,59) = 12.74, p < 0.0001) and N2 (F(2,59) = 12.74, p = 0.0002) latencies, compared to normal mice (Supplementary Figure 1Biv; Figure 8Fi–ii). Treatment with 5IIQ significantly increased P1 (F(2,47) = 24.17, p = 0.0054) and N2 (F(2,47) = 24.17, p = 0.0354) amplitudes with significantly decreased P1 latency (F(2,45) = 27.94, p < 0.0001), compared to vehicle-treated EAE mice. N2 latency did not differ significantly between 5IIQ- and vehicle-treated EAE mice.

Together, these results indicate that SARM1 inhibition with 5IIQ improves visual pathway function and facilitates partial recovery in EAE mice, despite not preventing structural RNFL thinning.

#### 5IIQ Treatment Selectively Dampens the Peripheral Immune Response During EAE

SARM1 activation is known to initiate axon damage in neurons through its NADase activity(Essuman et al., 2017; Gerdts et al., 2013; Osterloh et al., 2012). Although highly expressed in neurons, SARM1 is also expressed in other cell types(Doran et al., 2021; Gerdts et al., 2013; Jin et al., 2022; Osterloh et al., 2012; Sugisawa et al., 2024), and in human monocytes, SARM1 regulates pro-inflammatory cytokine expression during inflammation(Sugisawa et al., 2024). To determine how SARM1 inhibition via 5IIQ affects the peripheral immune response in EAE, splenocytes were isolated from WT mice at day 45, restimulated with MOG35–55 peptide ex vivo, and analyzed for cytokine and chemokine release (Figure 9). Consistent with past studies, stimulated splenocytes from EAE+V mice showed significantly increased pro-inflammatory cytokines, including TNF-α (t = 2.335, df = 17, p = 0.0399), IFN-γ (t = 37.12, df = 17, p = 0.0053), and IL-1α (t = 3.003, df = 17, p = 0.016), relative to CFA controls. Treatment with 5IIQ significantly decreased TNF-α (t = 2.983, df = 17, p = 0.0203) and IL-1α (t = 2.635, df = 17, p = 0.0346) compared to vehicle-treated mice; in contrast, IFN-γ and IL-1β remained significantly elevated in EAE+V relative to CFA controls but did not differ significantly between EAE+V and EAE+5IIQ groups. EAE induction also significantly increased IL-17 (t = 3.213, df = 10, p = 0.0013), CXCL1 (t = 5.959, df = 17, p = 0.005), and CXCL10 (t = 3.313, df = 17, p = 0.007) relative to CFA controls, and treatment with 5IIQ significantly decreased IL-17 (t = 1.972, df = 10, p = 0.0486), CXCL1 (t = 3.868, df = 17, p = 0.0025), and CXCL10 (t = 2.81, df = 17, p = 0.023) compared to vehicle-treated mice. EAE induction also significantly increased the anti-inflammatory cytokine IL-10 relative to CFA controls (t = 5.959, df = 10, p = 0.0020), and treatment with 5IIQ significantly decreased IL-10 release compared to vehicle-treated mice (t = 7.000, df = 10, p = 0.0011). Together, these findings indicate that 5IIQ treatment reduces axonal damage and selectively dampens a subset of the peripheral pro-inflammatory and chemokine response during EAE including TNF-α, IL-1α, IL-9, IL-10, IL-17, CXCL1, and CXCL10 while leaving other markers of systemic immune activation, including IFN-γ and IL-1β, unaffected.

**Figure 9:**
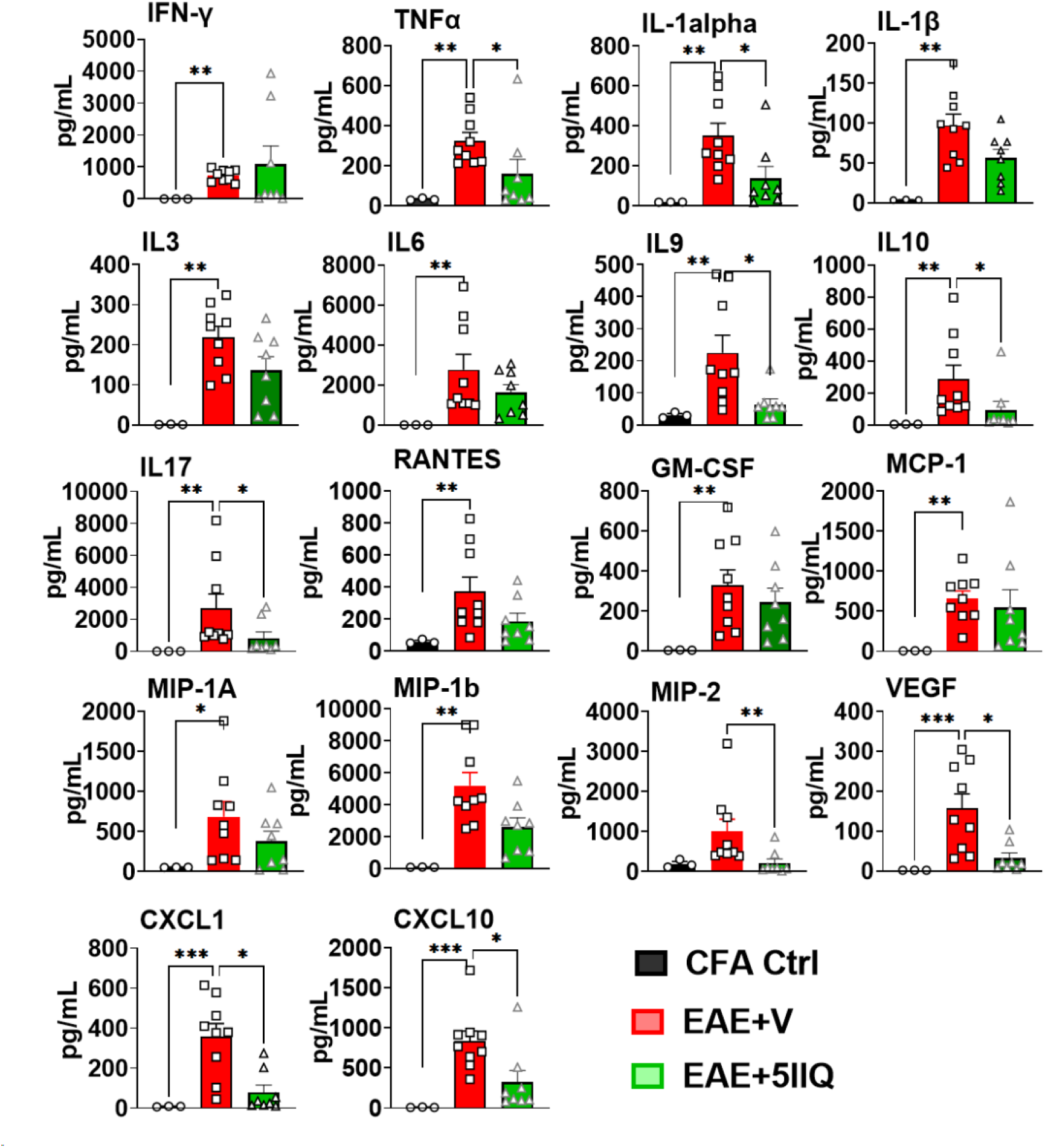
5IIQ treatment selectively dampens the EAE-induced peripheral cytokine and chemokine response in WT mice. Splenocytes from CFA control, EAE+vehicle (EAE+V), and EAE+5IIQ WT mice were restimulated ex vivo with MOG35–55 peptide, and culture supernatants were analyzed for cytokine and chemokine content. EAE induction significantly increased IFN-γ, TNF-α, IL-1α, IL-1β, IL-3, IL-6, IL-9, IL-10, IL-17, RANTES, GM-CSF, MCP-1, MIP-1α, MIP-1β, MIP-2, VEGF, CXCL1, and CXCL10 relative to CFA controls. Treatment with 5IIQ significantly reduced TNF-α, IL-1α, IL-9, IL-10, IL-17, VEGF, CXCL1, and CXCL10 relative to EAE+V, whereas IFN-γ, IL-1β, IL-3, IL-6, RANTES, GM-CSF, MCP-1, MIP-1α, and MIP-1β did not differ significantly between EAE+V and EAE+5IIQ. Data represents mean ± SEM (black, CFA Ctrl; red, EAE+V; green, EAE+5IIQ); n=6-10 mice/group, Unpaired t test with Welch’s correction, *p<0.05, **p<0.01, ***p<0.001, ****p<0.0001.

#### 5IIQ Treatment Protects RGCs and Optic Nerve Axons in EAE by Reducing SARM1 Expression, Independent of Effects on Myelination and microglia activation

To comprehensively evaluate the effect of 5IIQ treatment on retinal and optic nerve pathology in EAE, retinal cross sections and longitudinally sectioned optic nerves from normal, EAE+V, and EAE+5IIQ mice were assessed by IHC for markers of neuronal/axonal integrity, cell death, myelination, glial activation, and SARM1/NMNAT2 expression (Figure 10).

**Figure 10:**
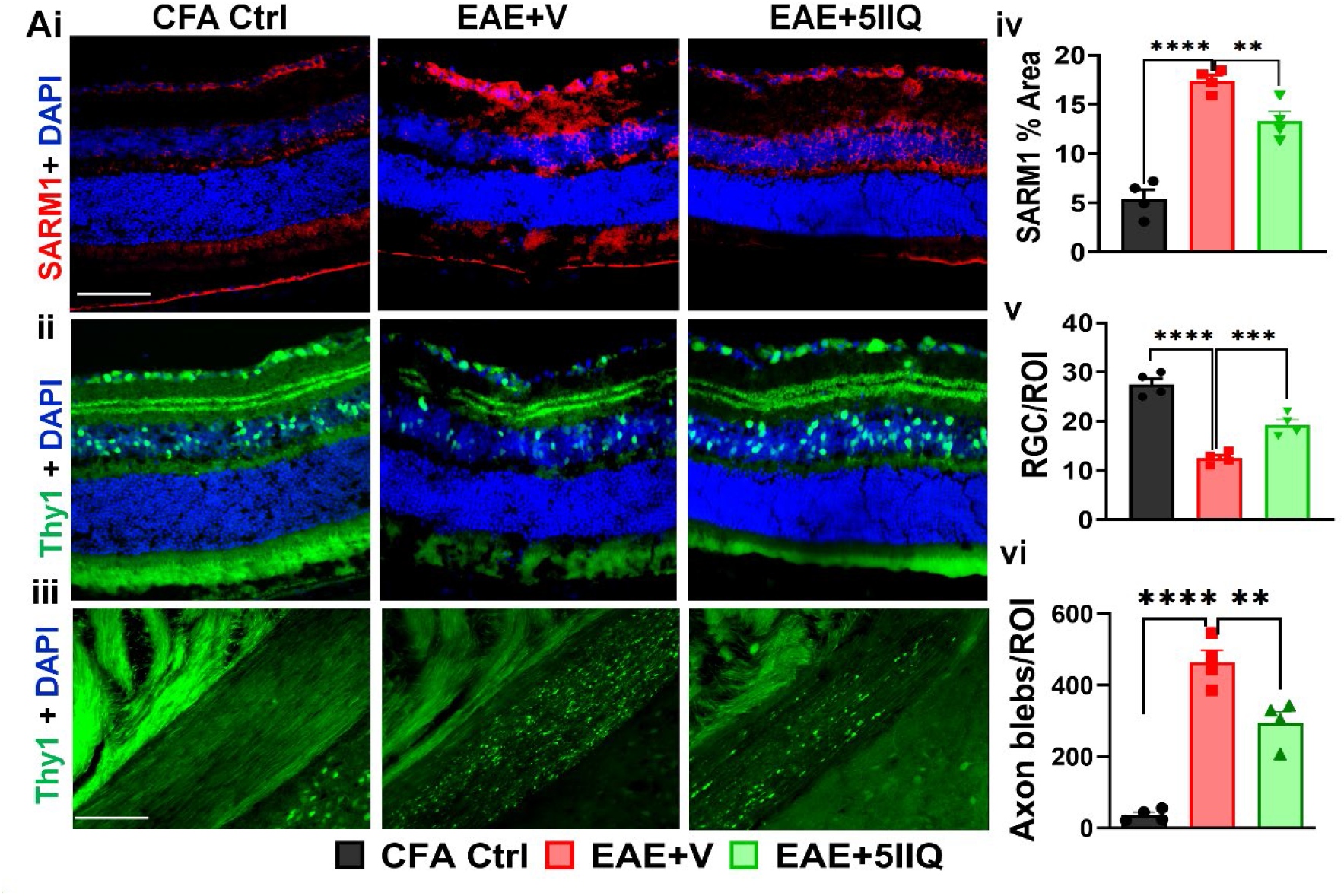
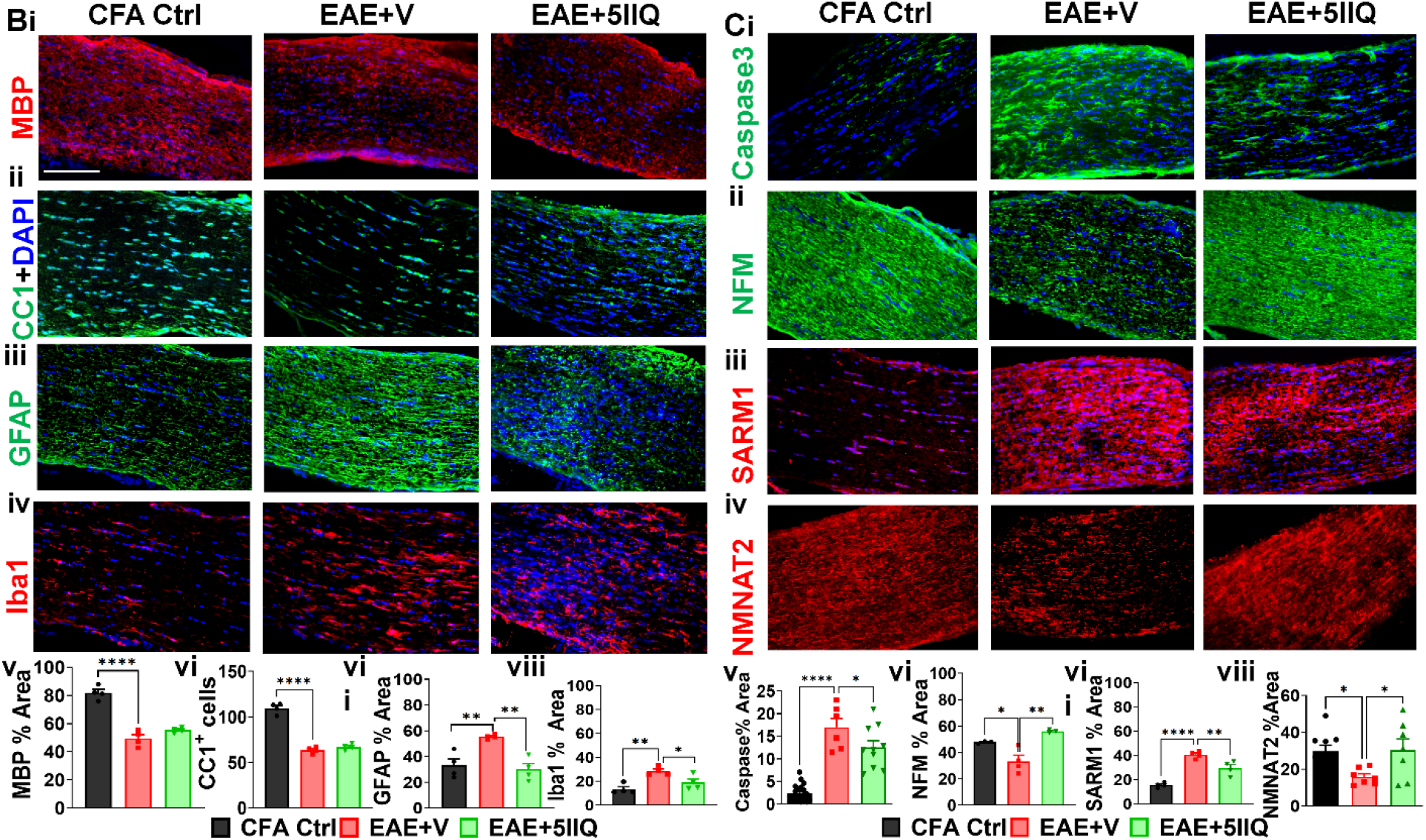
SARM1 inhibitor 5IIQ reduces retinal axon pathology despite unaltered clinical EAE scores. (Ai) Representative image showing retinal imaging location; retinal cross sections stained for SARM1 (red) show increased expression in EAE, reduced by 5IIQ treatment. Thy1-YFP positive RGCs and axons were assessed in retina (Aii, v) and optic tract (Aiii, vi): EAE significantly decreased Thy1-YFP+ RGCs, an effect attenuated by 5IIQ treatment. A significant increase in Thy1-YFP+ axon blebs was observed in the optic tract of EAE + vehicle mice, which were significantly reduced in 5IIQ treated groups. (B) Optic nerve sections were immunostained for MBP, CC1, GFAP, and Iba1: EAE decreased MBP and CC1 which were unaffected by 5IIQ treatment. The EAE + vehicle group showed an increased level of GFAP and Iba1. Only a decrease in GFAP was observed in 5IIQ-treated groups. (D) Optic nerve sections were immunostained for Caspase3, NFM, SARM1, and NMNAT2: EAE increased Caspase3 and SARM1 and decreased NFM and NMNAT2 relative to control; 5IIQ treatment reduced Caspase3 and SARM1 and restored NFM and NMNAT2 toward baseline. Data represent mean ± SEM (N=5–6 mice/group). One-way ANOVA with Bonferroni’s multiple comparisons test, *p<0.05, **p<0.01, ***p<0.001,****p<0.0001.

SARM1 immunostaining of Thy1-YFP retinal cross sections showed a significant increase in SARM1 expression across nearly all retinal layers in EAE+V mice compared to normal controls (F (2, 9) = 52.86, p < 0.0001), most prominently in the RNFL. Treatment with 5IIQ significantly decreased SARM1 immunoreactivity compared to vehicle-treated retinas (F (2, 9) = 52.86, p = 0.0142) (Figure 10Ai, iv). Consistent with this, quantification of Thy1-YFP⁺ RGCs showed a significant decrease in EAE+V mice compared to normal mice (F (2, 9) = 55.29, p < 0.0001), which was significantly preserved by 5IIQ treatment (F (2, 9) = 55.29, p = 0.0022) (Figure 10Aii, v). Because RGC loss is known to correlate with downstream optic nerve and optic tract pathology, we further examined Thy1-YFP⁺ axons in the optic tract, where EAE+V mice showed a significant increase in axonal blebbing F (2, 9) = 65.19, p <0.0001) that was significantly reduced by 5IIQ treatment (F (2, 9) = 65.19, p =0.0030) (Figure 10Aiii, vi). Together, these findings indicate that 5IIQ preserves RGCs and their axons throughout the visual pathway in association with decreased retinal SARM1 expression.

To determine whether 5IIQ’s protective effects extend to myelination and glial activation, optic nerve sections were stained for MBP, CC1 (mature OLs), GFAP (astrocytes), and Iba1 (microglia/macrophages) (Figure 10B). EAE+V optic nerves showed significant decreases in MBP (F (2, 9) = 61.03, p < 0.0001) and CC1 (F (2, 9) = 61.03, p < 0.0001), along with significant increases in GFAP (F (2, 9) = 12.57, p = 0.0023) and Iba1 (F (2, 9) = 10.67, p = 0.0026) immunoreactivity, compared to normal optic nerves. Treatment with 5IIQ did not alter MBP or CC1 expression compared to vehicle-treated optic nerves (Figure 10Bi, ii, v, vi), indicating that 5IIQ does not affect myelination or OL survival in EAE. In contrast, 5IIQ significantly reduced GFAP (F (2, 9) = 12.57, p=0.0026) and Iba-1 (F (2, 9) = 10.67, p = 0.0358) immunoreactivity compared to vehicle-treated optic nerves (Figure 10Biii, iv, vii, viii). These data indicate that 5IIQ selectively dampens astrocyte activation and microglial/macrophage reactivity. without affecting demyelination.

#### 5IIQ Reduces Cell Death and SARM1 Expression While Preserving Axonal Integrity and NMNAT2 in EAE Optic Nerves

To directly link SARM1/NMNAT2 signaling to axonal survival, optic nerve sections were immunostained for the apoptosis marker caspase-3, the axonal integrity marker NFM, and SARM1 and NMNAT2 (Figure 10C). EAE+V optic nerves showed significant increases in caspase-3 (F (2, 33) = 56.87, p < 0.0001) and SARM1 (F (2, 9) = 38.23, p <0.0001) expression, together with significant decreases in NFM (F (2, 7) = 12.84, p = 0.0337) and NMNAT2 (F (2, 20) = 4.054, p =0.0433) expression, compared to normal optic nerves (Figure 10Ci–viii). Treatment with 5IIQ significantly reduced caspase-3 expression (F (2, 33) = 56.87, p = 0.0317) and NFM loss (F (2, 7) = 12.84, p = 0.0033), and significantly decreased SARM1 (F (2, 9) = 38.23, p = 0.0081) while increasing NMNAT2 (F (2, 9) = 38.23, p = 0.0433) expression, compared to vehicle-treated EAE optic nerves.

Collectively (summarized in Supplementary Table 6), these findings demonstrate that 5IIQ treatment preserves RGCs and axonal integrity throughout the retina, optic nerve, and optic tract during EAE. Furthermore, this protection occurs in association with decreased SARM1 expression, restored NMNAT2 levels, reduced apoptosis, and reduced astrocyte activation, but independently of any effect on demyelination, OL survival, or microglial/macrophage reactivity. These results indicate that SARM1 inhibition confers neuroprotection in EAE through a mechanism that uncouples axonal and neuronal survival from the underlying inflammatory demyelinating disease process.

## Discussion

Axon damage remains the primary driver of irreversible disability in MS(Campbell and Mahad, 2018a). Inflammatory demyelination raises axonal energy demand beyond mitochondrial capacity, reversing the Na⁺/Ca²⁺ exchanger and triggering Ca²⁺-dependent calpain activation and cytoskeletal fragmentation(Stys, 1998). Approved MS therapies target the upstream inflammatory trigger; few directly protect axons(Albelo-Martinez and Rizvi, 2025). In this study, we used a combination of postmortem human tissue, genetic ablation, cell-type-specific gene manipulation, and pharmacological inhibition to establish SARM1 as a central, druggable driver of axonal degeneration in the visual pathway during CNS autoimmune demyelinating disease. Together, these complementary approaches converge on a consistent conclusion: SARM1 activity is upregulated at sites of axonal injury in both progressive MS tissue and EAE, and interrupting SARM1 signaling whether through germline knockout, RGC-restricted knockdown, or small-molecule inhibition preserves axonal and neuronal integrity without resolving the underlying immune-mediated inflammatory demyelination. This dissociation is, in our view, the most important and translationally relevant finding of the study, because it suggests that SARM1-targeted therapies could be layered onto existing immunomodulatory or novel remyelinating MS treatments as a neuroprotective adjunct rather than a substitute for them.

### SARM1 upregulation links human MS pathology to animal models

The elevated SARM1 immunoreactivity we observed in cerebellar white matter, the PC layer, the CA2 and SR regions of the hippocampus in progressive MS cases, co-localized with reduced NF200 staining, extends prior rodent-based descriptions of SARM1 biology to human neurodegenerative tissue. This finding provided the rationale for using EAE and ONC model(Cameron et al., 2020; Tang et al., 2011) as amenable systems to dissect SARM1’s contribution to axon loss, and the parallel elevation of SARM1 in EAE optic nerve alongside fragmented, beaded NFM staining and elevated serum NfL supports the validity of the visual pathway as a readout of SARM1-dependent axonal injury in this model.

### Germline SARM1 ablation reveals a structural, not immunological, protective mechanism

SARM1-/- mice subjected to EAE showed preserved NFM integrity(Viar K, 2020), fewer SMI-32+ axonal blebs, significant protection from demyelination and OL loss compared to WT EAE, yet showed similar clinical disease courses(Viar K, 2020), serum NfL elevations, and CD45+/CD3+ inflammatory infiltration that were largely indistinguishable from WT EAE animals. This pattern indicates that SARM1 loss acts downstream of, or in parallel to, the inflammatory cascade rather than by blunting it. It also implies that serum NfL, while a sensitive marker of neuroaxonal injury, may not fully capture the local structural protection conferred by SARM1 ablation, an important caveat for using NfL as a stand-alone biomarker of treatment response in future SARM1-targeted trials(Bosanac et al., 2021; Mani et al., 2025). The concurrent rise in NMNAT2 in SARM1KO optic nerve during EAE is consistent with a compensatory feedback loop within the same NAD+-homeostasis axis(Essuman et al., 2017; Gilley et al., 2015) and raises the possibility that combined SARM1 inhibition/NMNAT2 stabilization strategies could yield additive protection.

### SARM1 also shapes systemic innate immune tone independent of its axonal role

The novel finding that SARM1KO splenocytes exhibit an altered baseline cytokine/chemokine profile (elevated IL-1β, IL-1α, IL-9, RANTES, CXCL1, MIP-1α, MIP-2, and VEGF) and that several of these mediators show an inverted disease trajectory relative to WT animals, fits with SARM1’s known role as a negative regulator of MyD88-dependent TLR signaling(Carty et al., 2006). This suggests that germline SARM1 deletion produces two largely separable phenotypes: (1) a peripheral, disease-independent shift in innate immune set-point, and (2) a local, disease-dependent preservation of axons and myelin. Because these two phenotypes did not appear to interact in a straightforward manner (i.e., a “primed” innate immune system did not translate into worse or better clinical EAE), we favor a model in which the neuroprotective effect of SARM1 loss is cell-intrinsic to neurons/axons (SARM1 knockdown in nestin+ cells induced milder EAE (Liu et al., 2021a)) rather than secondary to altered systemic immunity a conclusion strengthened by the RGC-restricted knockdown data described below.

### Cell-autonomous manipulation of SARM1 in RGCs confirms a neuron-intrinsic protective mechanism

AAV2-mediated CRISPR knockdown of SARM1 in RGCs, using the contralateral eye as an internal control, produced partial preservation of RGC number, reduced SMI-32+ axonal blebbing, and reduced astrogliosis without significant modifications in ERG/VEP amplitude or latency in most comparisons. Conversely, RGC-restricted SARM1 overexpression worsened both functional (ERG/VEP) and structural outcomes relative to internal controls. The asymmetry between these two manipulations overexpression producing clear, internally controlled deficits, and knockdown producing more modest, partially inconsistent benefit is informative. We interpret this as reflecting the practical limits of a partial, local knockdown in the setting of an ongoing systemic autoimmune process: because EAE continuously drives new SARM1 expression and inflammatory injury throughout the disease course, incomplete local suppression of SARM1 may be outpaced by ongoing pathogenic signaling, whereas any residual or ectopic SARM1 activity from overexpression is sufficient to produce injury. This asymmetry mirrors the pharmacokinetic reality that will face any SARM1-directed therapy: near-complete and sustained target engagement is likely necessary for functional, and not just histological, benefit.

Prior work targeting SARM1 in RGCs shows a consistent split between protection limited to axons and protection extending to both axons and cell bodies. In the ONC model, both germline knockout and AAV-shRNA knockdown protect RGC axons but not the cell body, indicating an axon-specific degeneration pathway uncoupled from cell body death signaling(Fernandes et al., 2018). In contrast, SARM1 loss protects both axon and cell body in several inflammation and toxin-driven models such as rotenone-induced axon death(Finnegan et al., 2022), silicone oil-Induced ocular hypertension (SOHU) Glaucoma Model(Fang et al., 2023), and bead-induced ocular hypertension(Zhang et al., 2024). A TNFα-driven glaucoma model showed SARM1 downstream of both neuroinflammatory and necroptotic signaling leading to oligodendrocyte loss, axon degeneration, and RGC death(Ko et al., 2020). Most directly relevant to our results (Liu et al., 2023) compared germline knockout, retina ASO, and RGC-specific CRISPR knockdown across three different optic neuropathy models and found that none protected RGCs or optic nerve in EAE/optic neuritis, in contrast to robust protection in SOHU glaucoma and axon-only protection after traumatic injury. This convergence across independent methods indicates that EAE-driven optic neuropathy is resistant to RGC-restricted SARM1 loss specifically. Our findings that systemic 5IIQ achieved meaningful benefit in EAE suggest that SARM1 activity in surrounding cells, rather than the RGC itself, may need to be targeted to achieve protection in this disease context.

### Pharmacological SARM1 inhibition recapitulates and extends genetic findings and clarifies mechanism

The SARM1 inhibitors 5IIQ and CF3 protected primary cortical neurons from rotenone-induced, Complex-I-inhibition-driven axonal and dendritic loss, establishing efficacy in a metabolic stress paradigm relevant to mitochondrial dysfunction in neurodegeneration. In the ONC model, 5IIQ preserved NFM integrity and restored NMNAT2 expression without altering SARM1 expression itself or preventing RNFL thinning, a pattern that is mechanistically consistent with 5IIQ acting as a functional NADase inhibitor downstream of SARM1 induction, rather than as a suppressor of SARM1 transcription/translation(Hughes et al., 2021). Despite this dissociation between structural (RNFL) and molecular/functional outcomes, 5IIQ improved ERG and VEP amplitudes and latencies in ONC mice, indicating that a meaningful degree of functional rescue can occur even when overt RNFL thinning is not prevented. This is likely because retinal function depends on the health of surviving axons and synaptic circuitry rather than on RNFL thickness alone(Qiu et al., 2026).

In EAE, 5IIQ similarly failed to alter clinical disease score or RNFL thinning, but reduced serum NfL, improved ERG/VEP amplitude and latency, decreased optic nerve and retinal SARM1 expression, restored NMNAT2, reduced caspase-3^+^ apoptotic cells, and reduced astrogliosis, all without affecting demyelination, OL survival, or microglial/macrophage reactivity. This is essentially a pharmacological phenocopy of the SARM1KO phenotype with respect to inflammation and demyelination, but 5IIQ produced a different peripheral cytokine signature, selectively reducing TNF-α, IL-1α, IL-9, IL-10, IL-17, CXCL1, and CXCL10, while leaving IFN-γ and IL-1β both still significantly elevated in EAE relative to CFA controls unaffected, suggesting pharmacological inhibition and germline ablation are not immunologically equivalent. This is a key rationale for favoring a titratable, pharmacological approach.

### A convergent model: SARM1 uncouples axonal degeneration from CNS demyelinating inflammation

Across four independent manipulations germline knockout, RGC-specific knockdown, RGC-specific overexpression, and pharmacological inhibition the same qualitative pattern emerged: reducing SARM1 activity preserves neurofilament integrity, reduces axonal blebbing, limits RGC/neuronal loss, and often preserves visual function, while leaving clinical disease severity, T-cell/myeloid infiltration, and (except for the SARM1KO genotype) demyelination essentially unchanged. Consistent with 5IIQ acting on SARM1 activity rather than expression (see above), the reduction in SARM1 immunoreactivity with treatment in EAE likely reflects reduced injury-driven induction rather than a direct drug effect on transcription. This supports a model in which SARM1-dependent axonal self-destruction constitutes a distinct, targetable node located downstream of neuroinflammatory signals, yet but upstream of irreversible axon loss. This is analogous to the role SARM1 plays in Wallerian degeneration after mechanical injury but now shown to be operative in immune-mediated CNS injury as well.

### Several limitations qualify these conclusions

The human cohort was small and cross-sectional; germline knockout cannot rule out developmental compensation; AAV2-CRISPR knockdown was partial and mosaic; 5IIQ and CF3 are tool compounds whose selectivity was not directly tested; and discordance between functional (ERG/VEP) and structural (OCT/RNFL) outcomes indicates that RNFL alone is an insufficient surrogate for axon health. SARM1’s conserved roles in innate immunity and neurodevelopment also warrant caution regarding long-term inhibition. Further consideration relates to a recently described liability of orthosteric SARM1 inhibitors. Several groups have reported that base-exchange inhibitors (BEIs) which act as prodrugs converted by SARM1 into active-site-blocking NAD+ analogs paradoxically activate rather than suppress SARM1 at sub-inhibitory concentrations, worsening axonal degeneration both in vitro and in EAE in vivo (Leahey et al., 2025; Mani et al., 2025). This risk has been attributed to incomplete occupancy of the multiple adjacent active sites within the SARM1 TIR domain: when occupancy is partial, SARM1 can still oligomerize into a catalytically active state, accelerating rather than blocking NAD+ destruction (Leahey et al., 2025; Mani et al., 2025). Because 5IIQ acts through this same base-exchange mechanism(Bosanac et al., 2021; Feldman et al., 2022), it falls within the class of inhibitors for which this liability has been described, raising an important safety consideration for its therapeutic use. In our study, 5IIQ was administered at 25 mg/kg/day, a dose that produced consistent structural and functional axon protection without evidence of worsened outcomes in either the ONC or EAE model, arguing against paradoxical activation under this specific dose and regimen. However, we did not perform a full dose-response study, and we cannot exclude the possibility that lower, sub-therapeutic doses of 5IIQ, or prolonged treatment leading to declining tissue drug levels, could produce the same paradoxical activation reported for BEIs more broadly.

An alternative approach may sidestep this liability altogether: SARM1 inhibitors bearing electrophilic functional groups act instead through irreversible covalent modification of cysteine residues (C311 or C635) in the ARM autoregulatory or C-terminal TIR domains, providing durable allosteric inhibition that is not thought to be subject to low-dose paradoxical activation (Bosanac et al., 2021; Feldman et al., 2022). Future work should directly compare these covalent inhibitors against 5IIQ and other BEIs across a range of doses, both to establish formal dose–response relationships for 5IIQ and to confirm whether covalent, regulatory-domain-targeting inhibitors are indeed protected from this failure mode, as current evidence suggests (Bosanac et al., 2021; Feldman et al., 2022).

### Conclusions

Pharmacological SARM1 inhibition with 5IIQ provides structural and functional axonal protection in both a traumatic and a chronic inflammatory model of optic neuropathy, largely independent of demyelination or immune infiltration. Complementary genetic data reinforce SARM1 as a dose-sensitive, druggable node downstream of NMNAT2 depletion, while highlighting that only near-complete, tunable inhibition is likely to yield functional not just structural benefit. These findings support SARM1 inhibition as a promising neuroprotective strategy for MS, most likely as an adjunct to existing immunomodulatory and remyelinating therapies.

## Declaration of generative AI and AI-assisted technologies in the manuscript preparation process

During the preparation of this work, the author(s) used Claude (Anthropic) and Google Gemini pro to assist in editing and condensing some of the text and supplementary tables for clarity and conciseness. After using these tools, the author(s) reviewed, fact-checked, and rewrote as required. The author(s) take full responsibility for the accuracy and content of the publication work.

## Abbreviations

MS: multiple sclerosis, IHC: immunohistochemistry, EAE: experimental autoimmune encephalomyelitis, ONC: optic nerve crush, NfL: neurofilament, SARM1: Sterile alpha and TIR domain-containing protein 1, 5IIQ: 5-iodoisoquinoline. RGC: retinal ganglion cells, AAV2: Adeno-Associated Virus, serotype 2, SARM1 -/-: Sarm1 knockout

## Data availability statement

All raw data of IHC/VEP/ERG/OCT/Splenocyte cytokines is available on request.

