## Supplementary for "Breaking the link between demyelination and axon loss: SARM1 inhibition as a neuroprotective strategy for multiple sclerosis"

**Supplementary Figures**

**Supplementary Figure 1**: **Chronic MOG EAE has decreased RNFL and aberrant VEPs and ERGs.** (Ai–ii) EAE mice from Figure 2 show decreased RNFL as compared to CFA Controls. (Bi–vi) The EAE group also shows impaired visual function, with increased latency and decreased amplitude on ERG and VEP. N=6-8 mice/group, unpaired t test with Welch's correction. Data represents + SEM. *p<0.05, **p<0.01, ****p<0.0001.

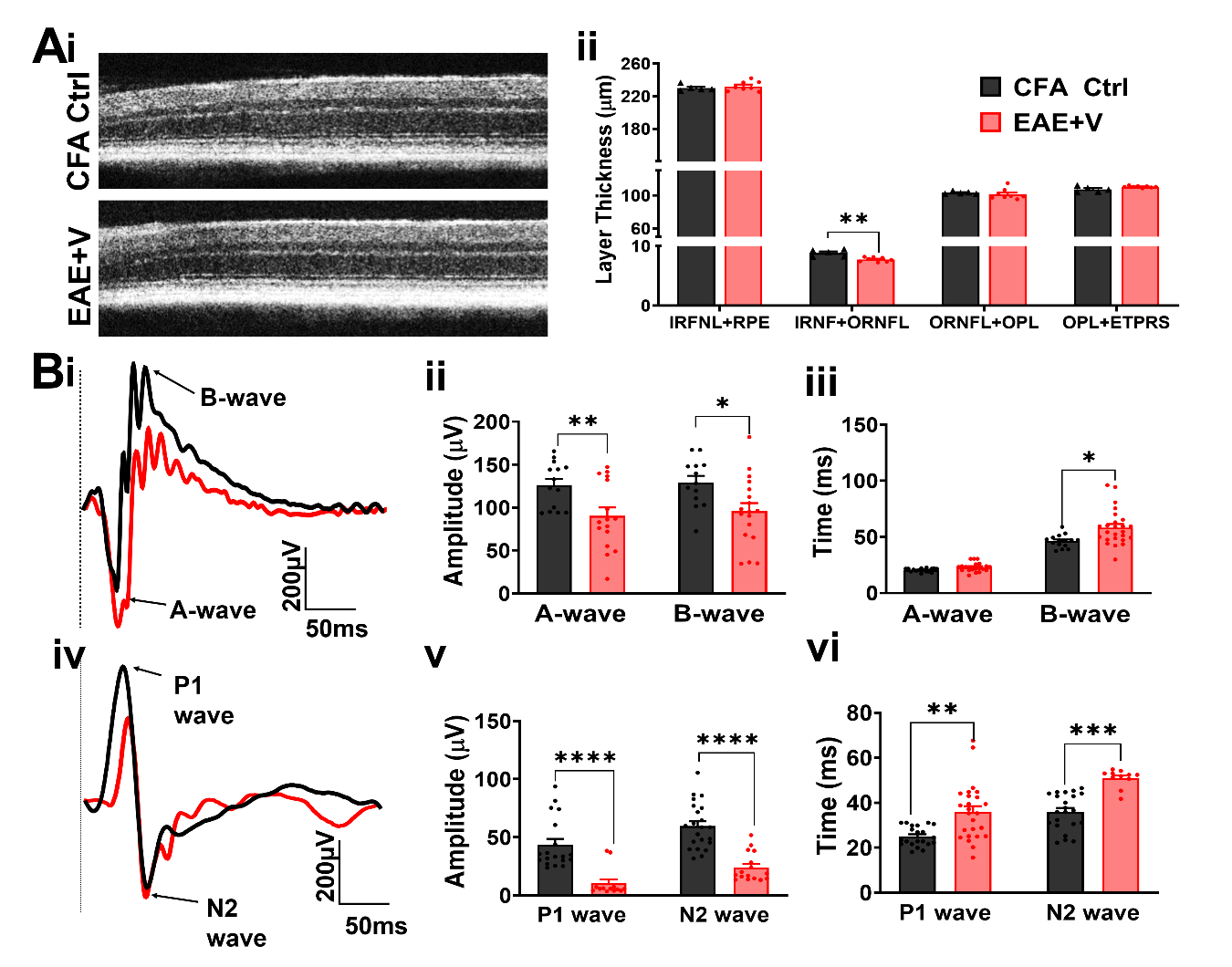

**Supplementary Figure 2**: **Feasibility of SARM1 inhibitors as potential therapeutic targets.** Axonal protection by SARM1 inhibitor: Primary mouse cortical neurons were treated with vehicle (DMSO+media) or rotenone for 6 hours followed by treatment for 5 days; No treatment, 10 uM 5-IIQ, 10 uM CF3, 1μM Rotenone (Rote); 1μM Rote+10uM 5-IIQ, 1μM Rote+10uM CF3. The cells were fixed and immunostained with MAP2 antibody (red), found in neuronal dendrites and perikaryal + neurofilament, NF200 (green), largely expressed in neuronal axons. +Rote alone treatment decimated neurons in culture. 5-IIQ and CF3 alone did not have an overall effect and looked like +vehicle cells. Significantly more intact neurons with less discontinuous MAP2 and NF200 staining were observed with 5-IIQ+Rote and CF3+Rote there were 3 different fields were imaged/slide. 3 separate slides/treatment. Mean + SEM. **p <0.01, ***p < 0.001, ****p<0.0001 by ordinary one‐way ANOVA with Bonferroni’s multiple comparison test.

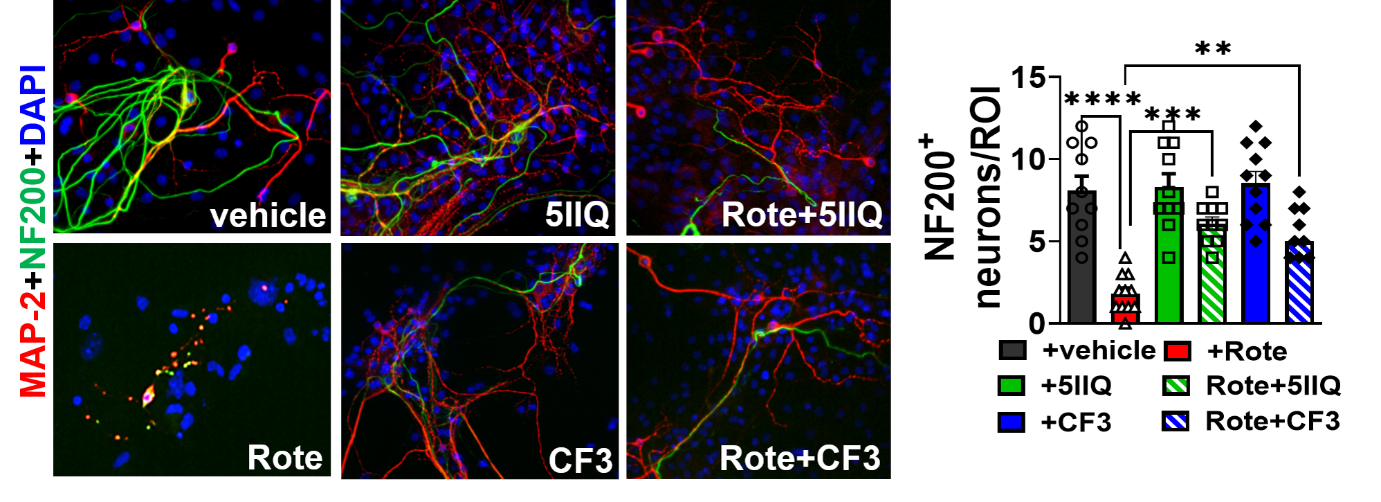

**Supplementary Figure 3:** **Optic nerve crush impairs axonal transport, RGC survival, retinal layer thickness, and visual function.** (A) Schematic of experimental design: CTB AlexaFluor 555 intravitreal injection, ONC injury 14 days later, and assessment of visual function (ERG, VEP) and ocular histology (retinal flat mounts, OCT) 14 days post-injury. (B) CTB Alexa 555 labeling of optic nerve sections shows reduced anterograde axonal transport in ONC relative to sham. (C, D) NeuN^+^/DAPI staining of retinal flat mounts and quantification shows a significant decrease in NeuN^+^ RGC area with ONC relative to sham. (Ei, ii) OCT imaging and layer thickness quantification show ONC significantly reduces INFL+ONFL thickness relative to sham, with no change in other retinal layers. (Fi–iii) Representative ERG traces show ONC reduces A- and B- wave amplitude relative to sham, with no change in latency. (Gi–iii) Representative VEP traces show ONC reduces P1 and N2 wave amplitude relative to sham, with no change in latency. N=6-8 mice/group, unpaired t test with Welch's correction. Data represents + SEM. *p<0.05, **p<0.01, ****p<0.0001.

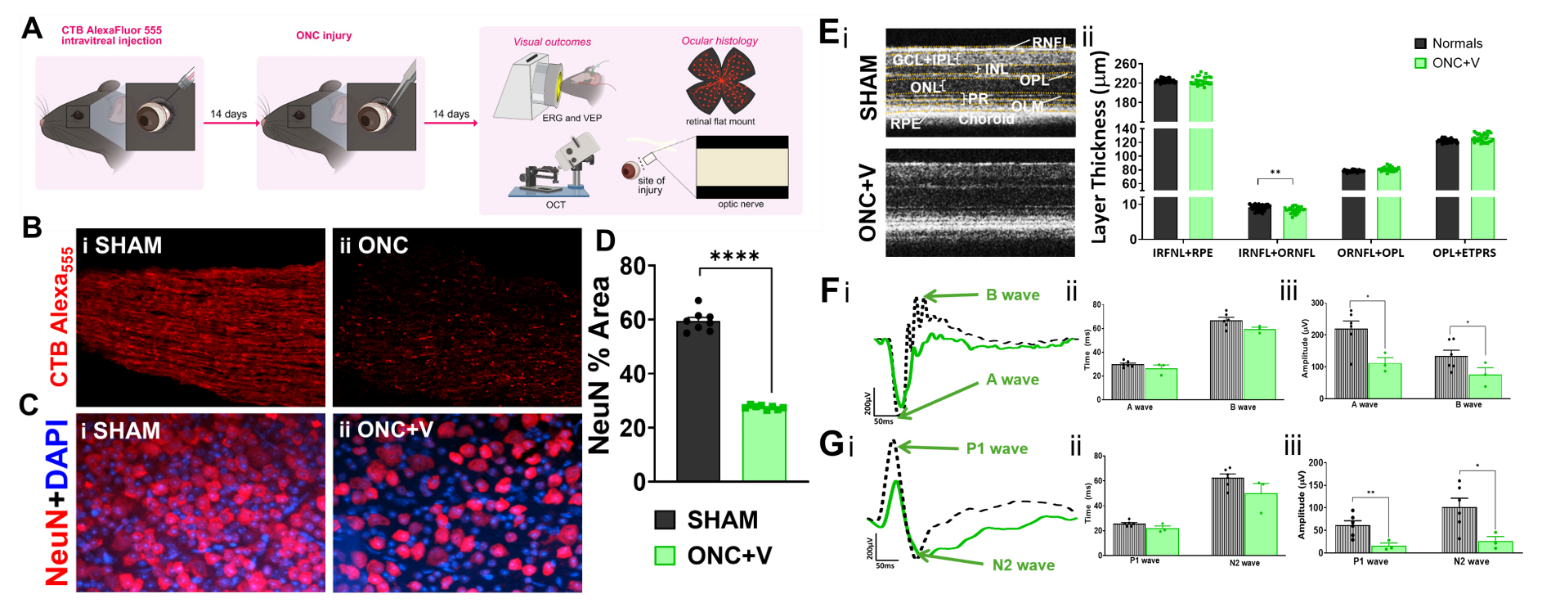

**Supplementary Tables**

| **Supplementary Table 1:** Details of disease and brain tissue pathology | | | | | |
| --- | --- | --- | --- | --- | --- |
|  | **ID** | **Age** | **Sex** | **Autolysis time (hrs)** | **Neuropathology** |
| **N** | HSB5214 | 61 | M | 19.5 | Unaffected Control |
|  | HSB5222 | 61 | M | 21.8 | Unaffected Control, micro-infarct (cerebrum), acute cortical microinfarct, frontal lobe (incidental) |
|  | HSB5293 | 41 | F | 11 | Normal, Liver failure |
|  | HSB4615 | 49 | M | 15 | Normal, CA, colon with metastasis to liver, depression, neuropath diagnoses: normal |
|  | HSB4130 | 67 | F | 11.8 | Normal, COPD, hypercholesterolemia, ypertension, macular degeneration, pulmonary emphysema, diabetes mellitus, depression, headache, anxiety, osteoporosis, nomal neuropath diagnosis |
|  | A250 | 79 | M | NA | Mild dementia |
|  | A040430 | 65 | M | NA | Pericardial hemorrhage, coronary atherosclerosis, hypertension, diabetes, chronic renal insufficiency |
|  | A16-13 | 57 | M | NA | Mild hypertensive vasulopathy, mild ventriculomegaly, incidental mineralization of hippocampal vessel |
| **MS** | HSB4212 | 50 | F | 18.9 | MS, hepatitis B, bed sore, chronic UTI |
|  | HSB5022 | 71 | M | 6 | MS |
|  | HSB5320 | 74 | F | 14.9 | MS, chronic UTI, neurogenic bladder, dysphagia, osteoarthritis, anemia, neuropathy, hypertension, depression (clinical), infarct: lacunar |
|  | HSB5025 | 73 | M | 2.9 | MS |
|  | HSB5024 | 67 | F | 11.8 | MS, hypertension, osteoporosis, hyperlipidemia |
|  | MS178 | 87 | M | 8 | MS |
|  | MS124 | 70 | F | 5 | PPMS |
|  | MS123 | 57 | M | 7 | RRMS |
|  | MS167 | 59 | F | 9.5 | PPMS |
| CA = cancer; COPD = chronic obstructive pulmonary disease; CVA = cerebrovascular accident; MS = multiple sclerosis; PPMS = primary progressive multiple sclerosis; RRMS = relapsing remitting multiple sclerosis; UTI = urinary tract infection | | | | | |

| **Supplementary Table 2: Antibody details** | | |
| --- | --- | --- |
| **Antibody** | **Manufacturer** | **Catalogue #** |
| MBP ^+^ | Abcam | AB40390 |
| CC1 ^+^ | GeneTex | GTX16794 |
| Olig2 ^+^ | Millipore | AB9610 |
| NFM ^+^ | Millipore | MAB1621 |
| SARM1 ^+^ | GeneTex | GTX77621 |
| CD45 ^+^ | Millipore | 05-1413 |
| GFAP ^+^ | Millipore | AB5541 |
| Iba1 ^+^ | Wako | 019-19741 |
| NMNAT2^+^ | ThermoFisher | H00023057-M04 |
| Caspase3^+^ | EMD Millipore | AM64 |
| NeuN^+^ | Invitrogen | MA5-33103 |
| NF200^#+^ | Sigma | N4142 |
| MAP2^#^ | Millipore | AB5622 |
| CD3^+^ | Cell signaling | E4T1B |
| SARM1^#^ | R&D Systems | MAB7037 |

| **Supplementary Table 3. Global SARM1 Knockout (SARM1KO) + EAE** | | | | |
| --- | --- | --- | --- | --- |
| **Category** | **Measure** | **WT EAE vs. WT CFA** | **SARM1KO EAE vs. SARM1KO CFA** | **SARM1KO EAE vs. WT EAE** |
| **Myelination** | MBP (optic nerve) | ↓ (F(3,16)=23.54, p<0.0001) | Small but significant ↓ (F(3,16)=23.54, p=0.0290) | SARM1KO EAE higher than WT EAE (F(3,16)=18.62, p=0.0079) |
| **Myelination** | Olig2 (OL transcription factor) | ↓ (F(3,15)=18.3, p<0.0001) | No significant difference vs. SARM1KO CFA | SARM1KO EAE higher than WT EAE (F(3,15)=18.3, p=0.0144) |
| **Myelination** | CC1 (mature OLs) | ↓ (F(3,19)=43.27, p<0.0001) | Small but significant ↓ (F(3,19)=43.27, p=0.0021) | SARM1KO EAE higher than WT EAE (F(3,19)=43.27, p<0.0001) |
| **Axon/neuronal damage** | Clinical EAE score | Standard EAE course | Earlier onset than WT EAE; comparable to WT by day 20 | Comparable by day 20 |
| **Axon/neuronal damage** | Serum NfL | ↑ (F(3,11)=84.22, p<0.0001) | ↑ (F(3,11)=84.22, p<0.0001) | No significant difference vs. WT EAE |
| **Axon/neuronal damage** | NFM (optic nerve) | ↓ (F(3,24)=4.816, p=0.0173) | No significant change vs. SARM1KO CFA | Preserved in SARM1KO EAE (structural protection) |
| **Axon/neuronal damage** | SMI-32+ axon blebbing | ↑ (F(3,19)=64.70, p<0.0001) | ↑ but attenuated (F(3,19)=64.70, p<0.0001) | SARM1KO EAE has significantly fewer blebs than WT EAE (F(3,19)=64.70, p<0.0001) |
| **Axon/neuronal damage** | NMNAT2 (optic nerve) | No significant change vs. WT CFA | ↑ vs. SARM1KO CFA (F(3,16)=8.671, p=0.0361) | — |
| **Immune/glial changes** | CD45 (microglia/macrophage) | ↑ (F(3,21)=13.53, p=0.0086) | ↑ (F(3,21)=13.53, p=0.0004) | Comparable increase to WT EAE |
| **Immune/glial changes** | CD3 (T cells) | ↑ (F(3,32)=27.15, p<0.0001) | ↑ (F(3,32)=27.15, p<0.0001) | Comparable increase to WT EAE |
| **Immune/glial changes** | Splenocyte baseline cytokines (IL-1β, IL-1α, IL-9, RANTES, CXCL1, MIP-1α, MIP-2, VEGF) | N/A (WT CFA baseline) | Elevated at baseline (SARM1KO CFA) vs. WT CFA | — |
| **Immune/glial changes** | Disease-associated trajectory: IL-9, RANTES, MIP-2, CXCL1 | Low at baseline, ↑ with EAE induction (expected pattern) | Elevated at baseline, ↓ with EAE induction (inverted trajectory) | Genotype-by-disease interaction; opposite direction from WT |

| **Supplementary Table 4. EAE + AAV2 (RGC-restricted SARM1 knockdown [KD] vs. overexpression [OE])** | | | |
| --- | --- | --- | --- |
| **Category** | **Measure** | **SARM1 KD vs. EAE/gRNA control** | **SARM1 OE vs. EAE/gRNA control** |
| **Axon/neuronal damage** | Clinical EAE score | Comparable across KD, OE, and control | Comparable across KD, OE, and control |
| Axon/neuronal damage | ERG A-wave amplitude | ↑ in both injected and internal control eyes (not locally specific) | ↓ specifically in injected eye vs. internal control (F(3,40)=14.67, p<0.0001) and vs. EAE/gRNA control (F(3,40)=14.67, p=0.0012) |
| Axon/neuronal damage | ERG B-wave amplitude | ↑ in both injected and internal control eyes (not locally specific) | No significant change reported |
| Axon/neuronal damage | ERG A/B-wave latency | No significant change (by ANOVA) | No significant change (by ANOVA) |
| Axon/neuronal damage | VEP P1 amplitude | No significant change (by ANOVA) | ↓ vs. EAE/gRNA control (F(5,41)=8.975, p=0.0224) |
| Axon/neuronal damage | VEP N2 amplitude | No significant change (by ANOVA) | ↓ vs. EAE/gRNA control (F(5,41)=8.975, p=0.0126) |
| Axon/neuronal damage | VEP P1 latency | No significant change | No significant change |
| Axon/neuronal damage | VEP N2 latency | No significant change | No significant change |
| **Axon/neuronal damage** | RGC survival (NeuN+) | Attenuated loss vs. OE/control (F(3,18)=17.65, p=0.0403) | Loss comparable to EAE control |
| **Axon/neuronal damage** | SMI-32+ axon blebbing | ↓ vs. EAE control/OE (F(3,20)=43.24, p=0.0001) | Comparable to EAE control |
| **Immune/glial changes** | GFAP (astrocytes) | ↓ vs. OE (F(3,22)=14.81, p=0.0130) | Higher than KD |
| **Immune/glial changes** | CD45 (microglia/macrophage) | ↑ vs. OE (F(3,21)=22.83, p=0.0034) | Lower than KD |
| **Immune/glial changes** | CD3 (T cells) | No significant difference vs. WT EAE or OE | No significant difference vs. WT EAE or KD |

| **Supplementary Table 5. ONC + 5IIQ (25 mg/kg/day)** | | | |
| --- | --- | --- | --- |
| **Category** | **Measure** | **ONC+Vehicle vs. Sham** | **ONC+5IIQ vs. Vehicle** |
| **Myelination** | MBP | No difference vs. sham | No difference vs. vehicle |
| **Myelination** | Iba1 (microglia/macrophage) | No difference vs. sham | No difference vs. vehicle |
| **Myelination** | GFAP (astrocytes) | No difference vs. sham | No difference vs. vehicle |
| **Axon/neuronal damage** | RGC survival (NeuN+) | ↓ (F (2, 21) = 167.2, p < 0.0001) | ↑ vs. ONC+V (F (2, 21) = 167.2, p < 0.0001) |
| **Axon/neuronal damage** | RNFL thickness (OCT) | ↓ (F (2, 80) = 10.20, p = 0.0001) | No significant difference vs. vehicle |
| **Axon/neuronal damage** | ERG A-wave amplitude | ↓ (t=2.839, df=7, p=0.0125) | ↑ (t=3.213, df=4, p=0.0163) |
| **Axon/neuronal damage** | ERG B-wave amplitude | ↓ (t=1.910, df=7, p=0.0489) | ↑ (t=3.273, df=4, p=0.0153) |
| **Axon/neuronal damage** | VEP P1 amplitude | ↓ (t=3.033, df=7, p=0.0095) | ↑ (t=2.842, df=4, p=0.0234) |
| **Axon/neuronal damage** | VEP N2 amplitude | ↓ (t=2.558, df=7, p=0.0188) | ↑ (t=2.882, df=4, p=0.0225) |
| **Axon/neuronal damage** | NFM (optic nerve) | ↓ (F (2, 21) = 11.74, p=0.0005) | ↑ / protected (F (2, 21) = 11.74, p=0.0015) |
| **Axon/neuronal damage** | SARM1 (optic nerve) | ↑ (F (2, 21) = 30.04, p<0.0001) | No significant change vs. vehicle |
| **Axon/neuronal damage** | NMNAT2 (optic nerve) | ↓ (F (2, 21) = 37.71, p<0.0001) | ↑ (F (2, 21) = 37.71, p<0.0001) |
| **Immune/glial changes** | Iba1, GFAP | Unchanged vs. sham (no gliosis in ONC) | No effect of 5IIQ |

| **Supplementary Table 6. EAE + 5IIQ (25 mg/kg/day)** | | | |
| --- | --- | --- | --- |
| **Category** | **Measure** | **EAE+Vehicle vs. Normal** | **EAE+5IIQ vs. Vehicle** |
| **Myelination** | MBP (optic nerve) | ↓ (t=8.299, df=5, p=0.0002) | No significant change |
| **Myelination** | CC1 (mature OLs) | ↓ (t=11.49, df=4, p=0.0002) | No significant change |
| **Axon/neuronal damage** | Clinical EAE score | Elevated vs. normal | No significant difference vs. vehicle |
| **Axon/neuronal damage** | Serum NfL | ↑ to 2,353 pg/mL (F (7, 35) = 5.448, p<0.0001) | ↓ to 1,393 pg/mL (F (7, 35) = 5.448, p = 0.0433) |
| **Axon/neuronal damage** | RNFL thickness (OCT) | ↓ (F (2, 18) = 3.879, p = 0.0397) | No significant difference vs. vehicle |
| **Axon/neuronal damage** | ERG A-wave amplitude | ↓ (F (2, 39) = 7.380, p = 0.0013) | ↑(F (2, 40) = 6.536, p = 0.0095) |
| **Axon/neuronal damage** | ERG B-wave amplitude | ↓ (F (2, 39) = 7.380, p = 0.0019) | No significant difference vs. vehicle |
| **Axon/neuronal damage** | ERG A-wave latency | ↑ (F (2, 39) = 7.380, p = 0.0030) | ↓ (F (2,41) = 6.329, p = 0.00352) |
| **Axon/neuronal damage** | ERG B-wave latency | ↑ (F (2, 39) = 7.380, p = 0.005) | ↓ (F (2, 41) = 7.380, p = 0.0068) |
| **Axon/neuronal damage** | VEP P1 amplitude/latency | ↓ amp (F (2, 47) = 24.17, p < 0.001); ↑ latency (F (2, 59) = 12.74, p < 0.0001) | ↑ amp (F (2, 47) = 24.17, p < 0.0054); ↓ latency (F (2, 45) = 27.94, p < 0.0001) |
| **Axon/neuronal damage** | VEP N2 amplitude/latency | ↓ amp (F (2, 47) = 24.17, p < 0.001); ↑ latency (F (2, 59) = 12.74, p < 0.0002) | ↑ amp (F (2, 47) = 24.17, p = 0.0354);No significant difference vs. vehicle |
| **Axon/neuronal damage** | SARM1 (retina) | ↑ (F (2, 9) = 52.86, p < 0.0001) | ↓ (F (2, 9) = 52.86, p = 0.0142) |
| **Axon/neuronal damage** | Thy1-YFP+ RGC survival | ↓ (F (2, 9) = 55.29, p<0.0001) | Preserved ↑ (F (2, 9) = 55.29, p = 0.0022) |
| **Axon/neuronal damage** | Optic tract axon blebbing | ↑ F (2, 9) = 65.19, p <0.0001) | ↓ (F (2, 9) = 65.19, p =0.0030) |
| **Axon/neuronal damage** | MBP and CC1 (optic nerve) | ↓ F (2, 9) = 61.03, p <0.0001) | No significant difference vs. vehicle |
| **Axon/neuronal damage** | NFM (optic nerve) | ↓ (F (2, 7) = 12.84, p=0.0337) | ↑ / protected (F (2, 7) = 12.84, p=0.0033) |
| **Axon/neuronal damage** | SARM1 (optic nerve) | ↑ ((F (2, 9) = 38.23, p<0.0001) | ↓ (F (2, 9) = 38.23, p = 0.0081) |
| **Axon/neuronal damage** | NMNAT2 (optic nerve) | ↓ (F (2, 20) = 4.054, p=0.0433) | ↑ (F (2, 20) = 4.054, p = 0.0433) |
| **Axon/neuronal damage** | Caspase-3 (apoptosis) | ↑ (F (2, 33) = 56.87, p < 0.0001) | ↓ (F (2, 33) = 56.87, p = 0.0317) |
| **Immune/glial changes** | GFAP (optic nerve) | ↑ (F (2, 9) = 12.57, p=0.0023) | ↓ (F (2, 9) = 12.57, p=0.0026) |
| **Immune/glial changes** | Iba1 (optic nerve) | ↑ (F (2, 9) = 10.67, p = 0.0026) | ↓ (F (2, 9) = 10.67, p = 0.0358) |
| **Immune/glial changes** | Splenocyte TNF-α | ↑ (t=2.335, df=17, p=0.0399) | ↓ (t=2.983, df=17, p=0.0203) |
| **Immune/glial changes** | Splenocyte IFN-γ | ↑ (t=37.12, df=17, p<0.0053) | No significant change |
| **Immune/glial changes** | Splenocyte IL-1α | ↑ (t=3003, df=17, p<0.016) | ↓ (t=2.635, df=17, p=0.0346) |
| **Immune/glial changes** | Splenocyte IL-1β | ↑ (t=3.993, df=10, p=0.0019) | No significant change |
| **Immune/glial changes** | Splenocyte IL-10 | ↑ (t=5.959, df=10, p=0.0020) | ↓ (t=7.000, df=10, p=0.0011) |
| **Immune/glial changes** | Splenocyte IL-17 | ↑ (t=3.213, df=10, p=0.0013) | ↓ (t=1.972, df=10, p=0.0486) |
| **Immune/glial changes** | Splenocyte CXCL1 | ↑ (t=5.959, df=17, p=0.005) | ↓ (t=3.868, df=17, p=0.0025) |
| **Immune/glial changes** | Splenocyte CXCl10 | ↑ (t=3.313, df=17, p=0.007) | ↓ (t=2.81, df=17, p=0.023) |
